# The dysregulated Pink1^-^ Drosophila mitochondrial proteome is partially corrected with exercise

**DOI:** 10.1101/2021.01.06.425659

**Authors:** Brad Ebanks, Thomas L Ingram, Gunjan Katyal, John R Ingram, Nicoleta Moisoi, Lisa Chakrabarti

## Abstract

One of the genes which has been linked to the onset of juvenile/early onset Parkinson’s disease (PD) is PINK1. There is evidence that supports the therapeutic potential of exercise in the alleviation of PD symptoms. It is possible that exercise may enhance synaptic plasticity, protect against neuro-inflammation and modulate L-Dopa regulated signalling pathways. We explored the effects of exercise on *Pink1* deficient *Drosophila melanogaster* which undergo neurodegeneration and muscle degeneration. We used a ‘power-tower’ type exercise platform to deliver exercise activity to *Pink1^-^* and age matched wild-type flies. Mitochondrial proteomic profiles responding to exercise were obtained. Of the 516 proteins identified, 105 proteins had different levels between *Pink1^-^* and wild-type (WT) non-exercised *D. melanogaster*. Gene ontology enrichment analysis and STRING network analysis highlighted proteins and pathways with altered expression within the mitochondrial proteome. Comparison of the *Pink1^-^* exercised proteome to WT proteomes showed that exercising the *Pink1^-^* flies caused their proteomic profile to return towards wild-type levels.

## Introduction

Parkinson’s disease (PD) is a progressive, irreversible neurodegenerative condition which affects over 6 million people across the globe(Ray Dorsey *et al*, 2018). The greatest risk factor for PD is advancing age, and with the number of people over the age of 60 expected to exceed 2 billion by the year 2050 (currently estimated to be over 900 million) there will soon be significantly higher numbers of people living with PD(Storey, 2018). The classical physical symptoms are known to be resting tremor, rigidity and bradykinesia. It is now also known that symptoms of PD include reduced quality of sleep as well as both cognitive impairments and poor mental health. In terms of the pathophysiology of the disease, the death of pigmented dopaminergic neurons in the substantia nigra pars compacta in PD patients is critical(Hirsch *et al*, 1988). Molecular characteristics of PD include the aggregation of α-synuclein leading to Lewy body formation, alongside mitochondrial dysfunction(Schulz-Schaeffer, 2010; Park *et al*, 2018).

Hereditary forms of PD can be either autosomal dominant or autosomal recessive, dependent upon the mutant gene involved. In both dominant and recessive forms of hereditary PD, autophagic and lysosomal pathways are both mechanistically implicated(Alegre-Abarrategui *et al*, 2009; Thomas *et al*, 2011; Zavodszky *et al*, 2014). One of the critical pathways which has been linked to the onset of juvenile/early onset PD is the PINK1/Parkin mitophagy pathway, a form of autophagy for the degradation of dysfunctional mitochondria. The role of mitophagy is to provide a quality control mechanism for the mitochondrial population within a cell, and this is a particularly crucial function in energetically demanding neuronal cells(Palikaras *et al*, 2018).

Mechanistically, PINK1 localizes with outer mitochondrial membrane (OMM) of depolarized mitochondria and then recruits and activates the E3 ubiquitin-ligase activity of Parkin via phosphorylation of its Ser65 residue (Shiba-Fukushima *et al*, 2012; Kondapalli *et al*, 2012). PINK1 has also been shown to phosphorylate the Ser65 residue of Ubiquitin, which aids in the activation of Parkin’s E3 ligase activity (Koyano *et al*, 2014; Kazlauskaite *et al*, 2014). Ultimately these events lead to the ubiquitination of OMM proteins by Parkin, the autophagic machinery then degrades the ubiquitinated mitochondria (Ordureau *et al*, 2014). Mutations in PINK1 (and Parkin) can result in autosomal recessive juvenile onset PD, with onset before the age of 40 years old (Cookson, 2012). While this form of PD is rare, a comprehensive understanding of it can improve the outcomes of patients with PINK1 mutations and also those with idiopathic PD due to their shared pathophysiology (Pankratz & Foroud, 2007). A recent publication by Sliter *et al*. reported that *Pink1* and *Parkin* mitigate STING induced inflammation, where both *Pink1^-/-^ and Prkn^-/-^* mice under exhaustive exercise have a strong inflammatory phenotype that is rescued by the concurrent loss of the STING pathway(Sliter *et al*, 2018). In the same year, Zhong *et al*. reported that newly synthesised oxidised mitochondrial DNA is exported to the cytosol and stimulates another of the innate immune responses, the NLRP3 inflammasome(Zhong *et al*, 2018).

Given the overlapping biology between PINK1 loss of function in PD and other forms of genetic and sporadic PD, PINK1 null mutants of animal models are hugely useful in studies of PD. *Pink1* loss of function *D. melanogaster* models were first developed in 2006, shortly after the first *parkin* loss of function mutant *D. melanogaster* models were being utilised in research(Park *et al*, 2006; Clark *et al*, 2006; Greene *et al*, 2003). These first studies found that *Pink1* loss of function resulted in mitochondrial dysfunction, compromised fertility in males, indirect flight muscle degeneration and associated locomotor defects, increased sensitivity to oxidative stress, and dopaminergic degeneration (Park *et al*., 2006). *Parkin* overexpression rescued many of the defects observed in the *Pink1* mutants, indicating the downstream function of Parkin in the now established PINK1/Parkin mitophagy pathway, reviewed here(Pickrell & Youle, 2015). More recent studies using *Pink1* mutant *D. melanogaster* have found that their neurons exhibit decreased levels of synaptic transmission, defective fission and reduced ATP levels due to decreased COXI and COXIV activity as well as non-motor symptoms such as learning and memory deficits, weakened circadian rhythms and electrophysiological changes in clock-neurons(Morais *et al*, 2009; Liu *et al*, 2011; Julienne *et al*, 2017).

It was first observed in 1992 that participation in exercise reduced the risk of the onset of PD in later years, while later data showed that this protection against PD risk is more obvious in males(Sasco *et al*, 1992; Yang *et al*, 2015). Many groups have presented data that show the therapeutic potential of exercise in the alleviation of patient symptoms(Ashburn *et al*, 2007; Goodwin *et al*, 2008; Murray *et al*, 2014). The biochemical mechanisms that would explain these observations are still unclear, but current evidence suggests that exercise may enhance synaptic plasticity, protect against neuroinflammation and modulate L-Dopa regulated signalling pathways(Shin *et al*, 2016; Jang *et al*, 2017; Klemann *et al*, 2018).

In this study we aimed to analyse the biochemical changes induced by exercise in the mitochondrial proteome of the *Pink1* loss of function mutant (*Pink1^-^*) *D. melanogaster*. As exercise is reported to both reduce the risk of onset and improve outcomes for Parkinson’s disease patients, we sought to characterise the biochemical changes that could underpin this improvement in our model of Parkinson’s disease. We focused on the mitochondria as their dysfunction is widely associated with Parkinson’s disease, and the *Pink1^-^* genetic model has a disrupted mitophagy pathway due to the absence of a functional PINK1 protein.

## Methods

### D. melanogaster stocks

Fly stocks were kindly provided to NM by Miguel Martins (MRC Toxicology Unit) and Alex Whitworth (MRC Mitochondrial Biology Unit). Fly stocks and crosses were maintained on standard cornmeal agar media at 25°C in a 12:12 light-dark cycle. The experiments were performed on males: WT (genotype *w1118)* and *Pink1^-^ (genotype Pink1B9/Y)*.

### D. melanogaster exercise

Approximately twenty wild type control (WT) or *Pi nk1^-^* flies, 1-4 days post-eclosion, were separated into glass vials filled with 5ml food. Exercised group vials were stoppered with cotton wool 6cm from the food; non-exercised group vials were stopped with cotton wool 1cm from the food, creating a physical barrier to activity. Both exercised and non-exercised groups were placed in racks on the ICE machine (**Appendix**) for 30 minutes per day for 7 days. The flies were exercised in the morning each day and were sacrificed by freezing at −80°C one hour after the final exercise bout. Comparison groups were exercised and non-exercised WT and *Pink1^-^ D. melanogaster*.

### Mitochondrial isolation

Groups of twenty WT or *Pink1^-^ D. melanogaster* were homogenised in 100-200µl mitochondrial extraction buffer (50mM Tris-Cl pH 7.4, 100mM KCL, 1.5mM MgCl2, 1mM EGTA, 50mM HEPES and 100mM sucrose) by 5 minutes of manual homogenisation using a 1.2-2ml Eppendorf micro-pestle (Sigma-Aldrich). The homogenate was centrifuged at 800g for 10 minutes, at 4°C, to remove the insoluble fraction. Supernatants from the first centrifugation were centrifuged at 1,000g for 10 minutes at 4°C to pellet the nuclear fraction. Supernatants from the second centrifugation were centrifuged at 13,200g for 30 minutes at 4°C to pellet the mitochondrial fraction. The protein content was determined by Bradford assay (µg/µl) and mitochondrial fractions were stored at −80°C.

### 2D-gel electrophoresis

50µg of the mitochondrial fraction were added to rehydration solution (8M urea, 2% CHAPS, 2% IPG Buffer, 0.1% bromophenol blue). 20mM DTT was added to an aliquot of rehydration solution directly before use. The standard protocol according to manufacturer instructions was followed(Life Technologies, 2012). Briefly, sample was applied to rehydrate ZOOM IPG strips for an hour at room temperature followed by iso-electric focussing using the ZOOM IPG (Life Technologies) system and pH 3-10 (non-linear) ZOOM IPG strips. Gels were stained (SimplyBlue™ SafeStain, Life Technologies) and imaged (ImageQuant 300, GE Healthcare Life Sciences). Analyses were performed using SameSpots software (Totallab) (one-way ANOVA). Three pooled biological replicates were included for each of the four groups.

Samples were analysed by the Centre of Excellence in Mass Spectrometry at University of York(Centre for Excellence in Mass Spectrometry, 2019). Briefly, proteins were reduced and alkylated, followed by digestion in-gel with trypsin. Matrix Assisted Laser Desorption Ionization Tandem Time-of-Flight mass spectrometry (MALDI-TOF/MS) was used to analyse the samples. The generated tandem MS data was compared against the NCBI database using the MASCOT search programme to identify the proteins. De novo sequence interpretation for individual peptides were inferred from peptide matches.

### Label-free proteomics

30µg/µl of each mitochondrial fraction was prepared with 4X LDS sample buffer and 4mM DTT. Samples were run in triplicate on a 4-12% Bis-Tris gel in 1X MES SDS running buffer for 40 minutes at 200V (all Invitrogen). Three biological replicates were run for each of the four groups. Individual gel lanes were excised and placed into separate Eppendorf tubes.

Samples were analysed by the Centre of Excellence in Mass Spectrometry at University of York(Centre for Excellence in Mass Spectrometry, 2019). Briefly, protein was in-gel digested post-reduction and alkylation. The resulting extracted peptides were analysed over 1-hour LC-MS acquisitions with elution from a 50cm, C18 PepMap column onto a Thermo Orbitrap Fusion Tribrid mass spectrometer using a Waters mClass UPLC. Extracted tandem mass spectra were searched against the combined drosophila melanogaster and Saccharomyces cerevisiae subsets of the UniProt database. Protein identifications were filtered to achieve <1% false discovery rate as assessed against a reverse database. Identifications were further filtered to require a minimum of two unique peptides per protein group.

For relative label-free quantification, extracted ion chromatograms for identified peptides were extracted and integrated for all samples. A maximum mass deviation of 3 ppm and retention time drift of 3 mins were set. Resulting quantifications were further filtered to an arbitrary PEAKS quality factor of 5 for feature mapping and required a minimum of two aligned features from a minimum of two unique peptides per protein quantification. Protein abundances were normalised between samples based on total identified peptide ion area.

### Gene Ontology enrichment analysis

gProfiler was used to undertake Gene Ontology (GO) enrichment analysis for the label-free mass spectrometry identified proteins with significant expression differences between each of the four experimental groups(Raudvere *et al*, 2019). KEGG, Molecular Function (MF), Biological Process (BP) and Cellular Compartment (CC) enrichment analyses are generated by gProfiler are presented.

### Protein-protein interaction network analysis

Differences in expression of proteins between groups were further analysed using the STRING database v.11.0(Szklarczyk *et al*, 2019). The platform was used to create protein-protein interaction (PPI) networks based upon the differentially expressed proteins (DEPs) observed between groups.

## Results & Discussion

We subjected male *Pink1* loss of function mutant (*Pink1^-^*) and WT *D. melanogaster* to a seven-day exercise regimen, whilst maintaining groups of unexercised *Pink1^-^* and WT as controls. The mitochondria of the four groups were then isolated and investigated using 2D gel electrophoresis and label-free mass spectrometry analyses to determine changes in their mitochondrial proteome. The 2D gel electrophoresis method allowed for a fast and simple separation of proteins, which served as a scoping method to identify some of the most significant changes in expression. Label-free mass spectrometry analyses generated proteomes that we used in network enrichment analysis. Previously we have used proteomic profiling to characterise mitochondrial populations in both mice and long-lived pipistrelle bats, and here we apply this to better understand the PD phenotype as well as possible changes due to exercise(Pollard *et al*, 2016, 2019).

### 2DE-MS identified a general reduction in protein expression with exercise in the *Pink1* mutant flies

We isolated mitochondria from exercised *Pink1^-^ D. melanogaster* and performed 2DE-MS on these fractions. All proteins identified as changed with our exercise intervention were reduced in expression (**Table 1**). PINK1 is recognised as having a central role in mitophagy, ensuring a healthy pool of mitochondria are maintained(Clark *et al*, 2006; Park *et al*, 2006). Dysfunctional mitophagy results in an accumulation of damaged and dysfunctional mitochondria and their contents(Anzell *et al*, 2018).

**Table 1.** Expression changes between exercised Pink1-flies and non-exercised Pink1-flies. Changes in expression were determined by 2DE-MS.

| **Protein identity** | **ANOVA (p)** | **Fold Change** | **Exercise related change** |
| --- | --- | --- | --- |
| Tropomyosin-1, isoforms 33/34 | 0.022 | 1.4 | ↓ |
| Tropomyosin-2 |  |  |  |
| Acyl-coenzyme A dehydrogenase | 0.015 | 1.4 | ↓ |
| Isocitrate dehydrogenase | 0.008 | 1.4 | ↓ |
| Enolase | 0.022 | 1.3 | ↓ |
| Probable isocitrate dehydrogenase [NAD] subunit alpha | 0.015 | 1.5 | ↓ |
| Glycerol-3-phosphate dehydrogenase [NAD(+)] |  |  |  |
| Pyruvate dehydrogenase E1 component subunit beta |  |  |  |
| Pyruvate dehydrogenase E1 component subunit beta | 0.048 | 1.3 | ↓ |
| Aldo-keto reductase, isoform C |  |  |  |
| Alcohol dehydrogenase | 0.024 | 1.3 | ↓ |
| CG9992, isoform A |  |  |  |

It is possible that alternate mitophagy pathways can compensate for PINK1 and are upregulated during exercise, this could account for the sweeping reductions in protein levels observed here. Indeed, alternative proteins have been identified which promote PINK1/PARKIN-independent mitophagy: AMBRA1, HUWE1 and IKKα(Di Rita *et al*, 2018). However, it could also be suggested that the energetic demands of the exercise result in less energy available to protein synthesis pathways. In this instance, exercise would likely affect proteostasis more broadly, beyond just the mitochondrial proteome.

We proceeded to pursue the directionally homologous 2DE-MS results by obtaining a global topology of the mitochondrial protein changes that occur to *Pink1^-^ D. melanogaster* due to exercise intervention using a label-free proteomics method.

### GO annotation of identified label-free proteins and proportion identified that are localised to mitochondria

Non-gel-based label-free proteomic analyses identified 516 proteins from the mitochondrial fractions of *Pi nk1^-^* and WT *D. melanogaster* (**Supp. table 1**). GO and KEGG analyses showed that these fractions were enriched for mitochondrial processes and pathways, confirming the efficacy of our fractionation methodology (**Fig. 1**). The top term of the GO cellular compartment analysis was cytoplasm, followed by mitochondrion and sub-mitochondrial compartments, many mitochondrial proteins are known to also localise to the cytoplasm(Yogev & Pines, 2011; Ben-Menachem *et al*, 2011).

**Figure 1.**
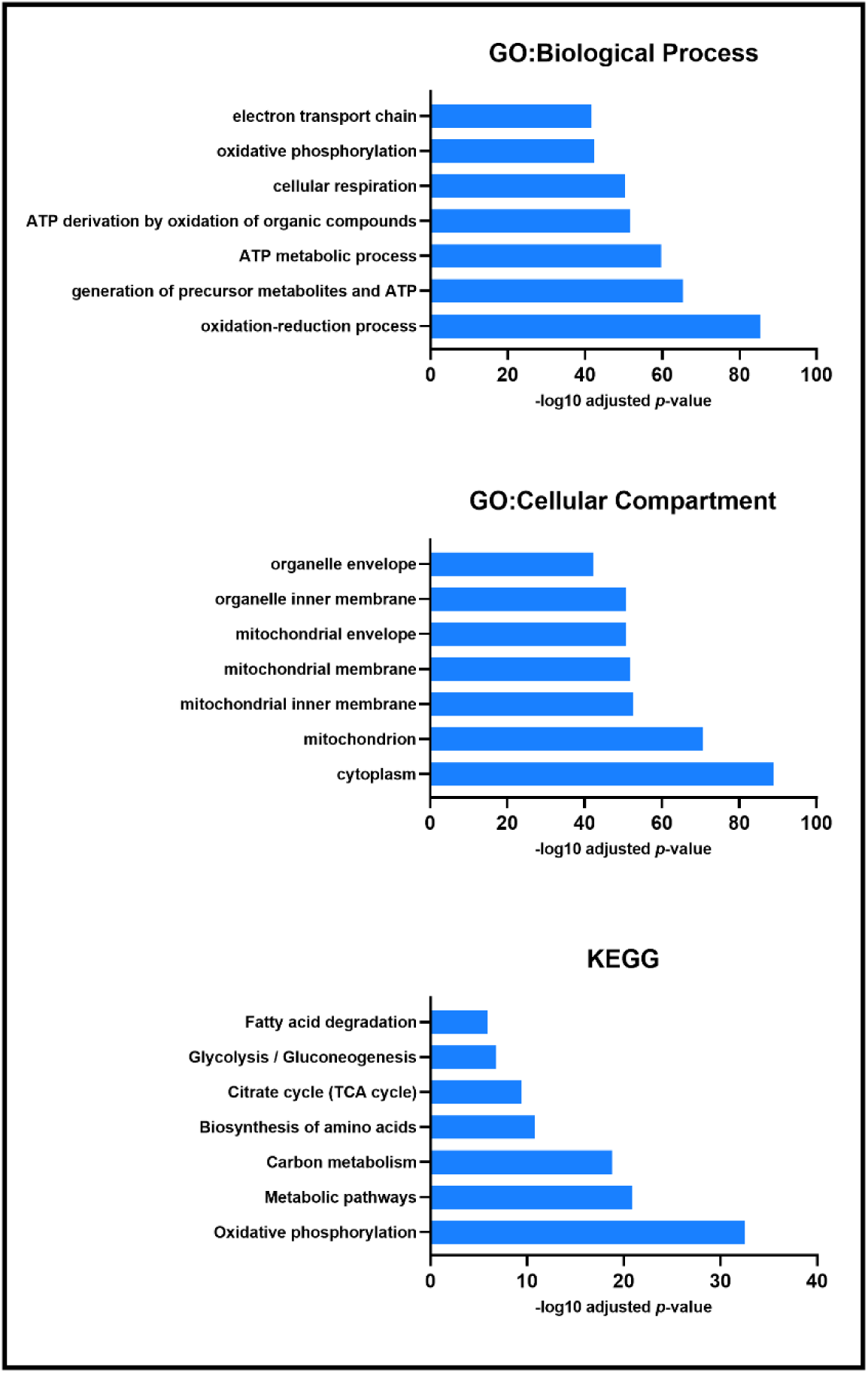
Enrichment analysis of all proteins identified by label-free proteomics. Biological process, cellular compartment and KEGG enrichment analysis each presented processes associated with mitochondrial function and physiology.

### *Pink1^-^* flies have decreased levels of proteins in energy metabolism pathways

Label-free proteomics highlighted 105 differently expressed proteins between non-exercised WT and *Pink1^-^* groups (**Supp. Table 3**). Ten proteins were shown to be reduced in expression in *Pink1^-^* compared to WT. We found that *Pink1^-^* flies have reductions in mitochondrial processes associated with energy metabolism, with the top GO biological process terms being oxidative phosphorylation, electron transport chain, ATP metabolic process and oxidation-reduction process (**Fig 2**). The deficiencies in mitochondrial oxidative phosphorylation, the electron transport chain and specifically in the activity of Complex I in Parkinson’s disease are well established, and this aligns with our findings from the GO analysis of the proteomics data(Parker *et al*, 1989; Schapira *et al*, 1990; Shoffner *et al*, 1991; Finsterer *et al*, 2001). Specific subunits of complexes within the electron transport chain that decreased in expression include NADH dehydrogenase (ubiquinone) 75 kDa subunit isoform B (complex I), GH01077p (complex III in *D.* melanogaster), HDC00331 (complex IV in *D. melanogaster*) and Levy isoform A (complex IV in *D. melanogaster*). While complex I and complex IV have been reported as dysfunctional in PD, reduced expression or decreased activity for complex III isn’t well documented(Keeney *et al*, 2006; Müftüoglu *et al*, 2004; Schapira *et al*, 1990). However, decreased Complex II/III activity has been shown in platelets of untreated Parkinson’s disease patients(Haas *et al*, 1995).

**Figure 2.**
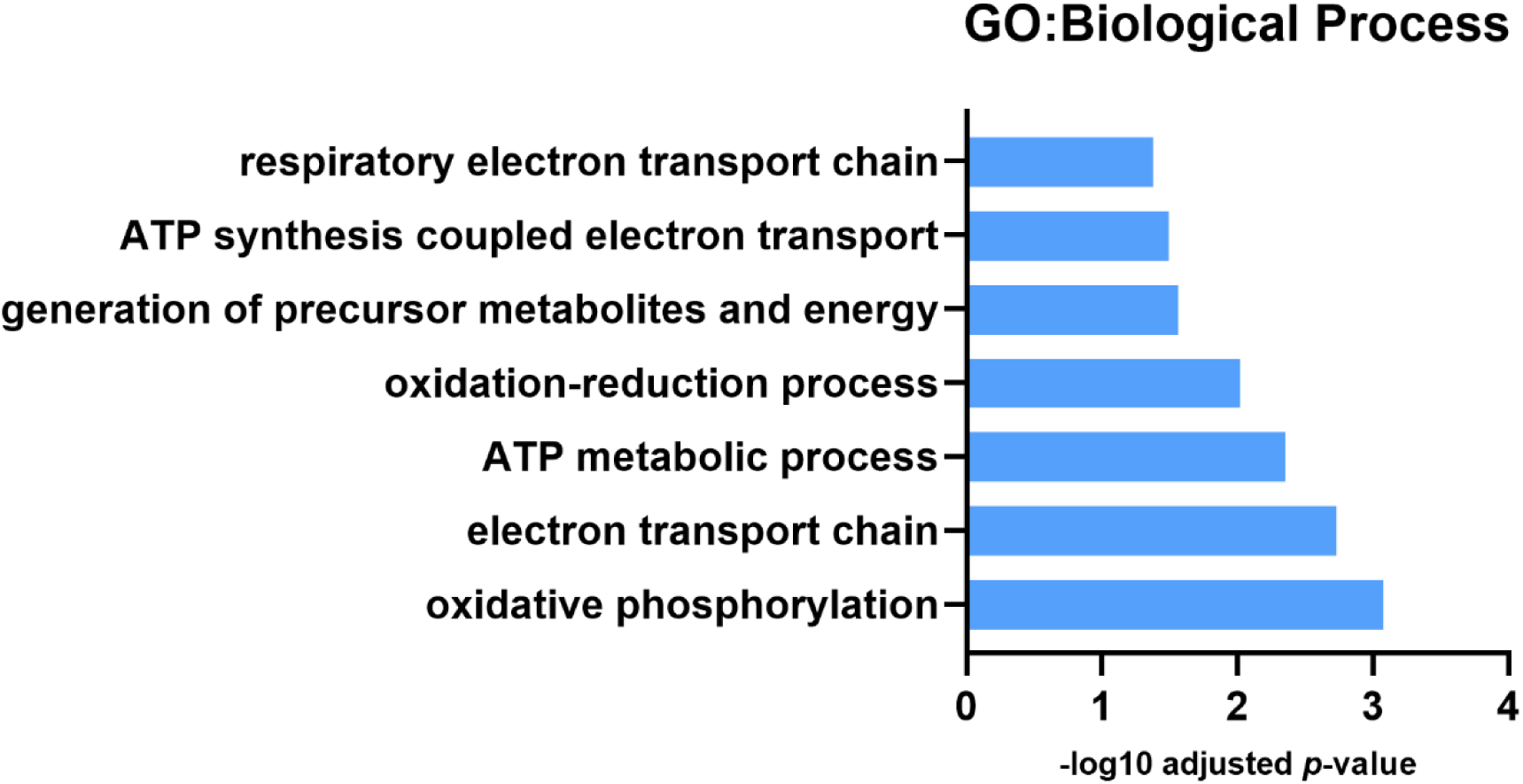
GO: Biological Process analysis for downregulated protein expression differences between Pink1^-^ non-exercised and wild-type non-exercised flies. Pink1^-^ flies have reduced expression of proteins involved in mitochondrial respiration and oxidative phosphorylation.

It is interesting to note that most (95/105) of the differentially expressed proteins were more highly expressed in *Pink1^-^ D. melanogaster*. GO biological process analysis showed these proteins to be enriched for redox processes, cytoplasmic translation, cellular amide metabolic processes and fatty acid derivative biosynthetic processes. GO cellular compartment analysis showed that the more highly expressed proteins in *Pink1^-^* were enriched for cytoplasmic ribosomes, organelle membranes and endoplasmic reticulum. KEGG pathway analysis paralleled these findings, highlighting the identified proteins as involved in fatty acid metabolism, ribosomes and one carbon pool by folate.

There is evidence linking fatty acid metabolism and function to Parkinson’s disease, with proteins identified by GWAS studies, supressed β-oxidation, and physical interaction between α-synuclein and fatty acids potential being key factors(Chang *et al*, 2017; Li *et al*, 2018; Saiki *et al*, 2017; Sharon *et al*, 2001; Karube *et al*, 2008). Early studies into the effect of α-synuclein (*SNCA*) gene deletion on lipid metabolism in mice reported reduced palmate uptake and altered palmate metabolism in the brain, reduced acyl-CoA Synthetase activity that resulted in reduced arachidonic acid uptake and turnover, and increased docosahexaenoic acid brain mass, incorporation and turnover(Golovko *et al*, 2005, 2006, 2007). Our own work shows differences in arachidonic acid derivatives in Parkinson’s disease mitochondria(Ingram *et al*, 2020).

The data presented here show an enrichment of the folate metabolic pathway, not reported previously. It may be that in *Pink1^-^* flies this is a compensatory mechanism. KEGG analysis highlighted the metabolism of folate (vitamin B_9_) as enriched in *Pink1^-^ D. melanogaster*. B-vitamins, in particular folate, are well studied in the context of Parkinson’s disease due to the observation of homocysteine (a methionine cycle metabolite) having neurotoxic effects(Lipton *et al*, 1997; Müller *et al*, 1999; Duan *et al*, 2002; Martignoni *et al*, 2007). It is hypothesised that the administration of B-vitamins can drive the synthesis of methionine, thus reducing intracellular homocysteine(Mayer *et al*, 2002; Maron & Loscalzo, 2009; Shen, 2015). However the relationship between B -vitamins, neurotoxicity and Parkinson’s disease is complex and a consensus has yet to be established. Some data show either little correlation between homocysteine levels and B vitamins including B6, folate and B12 while others show contradictory results, including elevated homocysteine levels and decreased folate levels in Parkinson’s disease patients(Xie *et al*, 2017; Shen *et al*, 2019; Dong & Wu, 2020; Shen, 2015).

### Exercise reduces measured protein levels in *Pink1^-^* flies

2DE analysis of *Pink1^-^ D. melanogaster* revealed reductions in twelve proteins with exercise (**Table 1**). Label-free proteomics showed a similar pattern in *Pink1^-^* exercised *D. melanogaster*; of the fifty-seven protein differences, fifty-five were reductions of protein expression with exercise (**Supp. table 2**). GO:MF and KEGG pathway analysis determined that the terms ribosomal and fatty acid metabolism were significantly represented in the proteins with reduced expression (**Fig 3**). Interestingly, these terms were shown to be increased in *Pink1^-^* compared to WT flies that had not been exercised. This suggests that in *Pink1^-^ D. melanogaster* exercise returns levels of protein expression towards WT values. It has been shown that PINK1 interacts with the protein translation pathway and that increased protein translation in *Pink1^-^ D. melanogaster* causes an exacerbated *Pink1^-^* phenotype. Taking this a step further, it was shown that 40S ribosomal subunit S6 (RpS6) RNAi was able to mitigate the *Pink1^-^* phenotype(Liu & Lu, 2010). These data suggest improper protein translation regulation is involved in the pathogenesis of PD and that inhibition of this pathway mitigates progression. Exercise appears to be able to reverse the upregulated protein translation pathways found in *Pink1^-^ D. melanogaster*.

**Figure 3.**
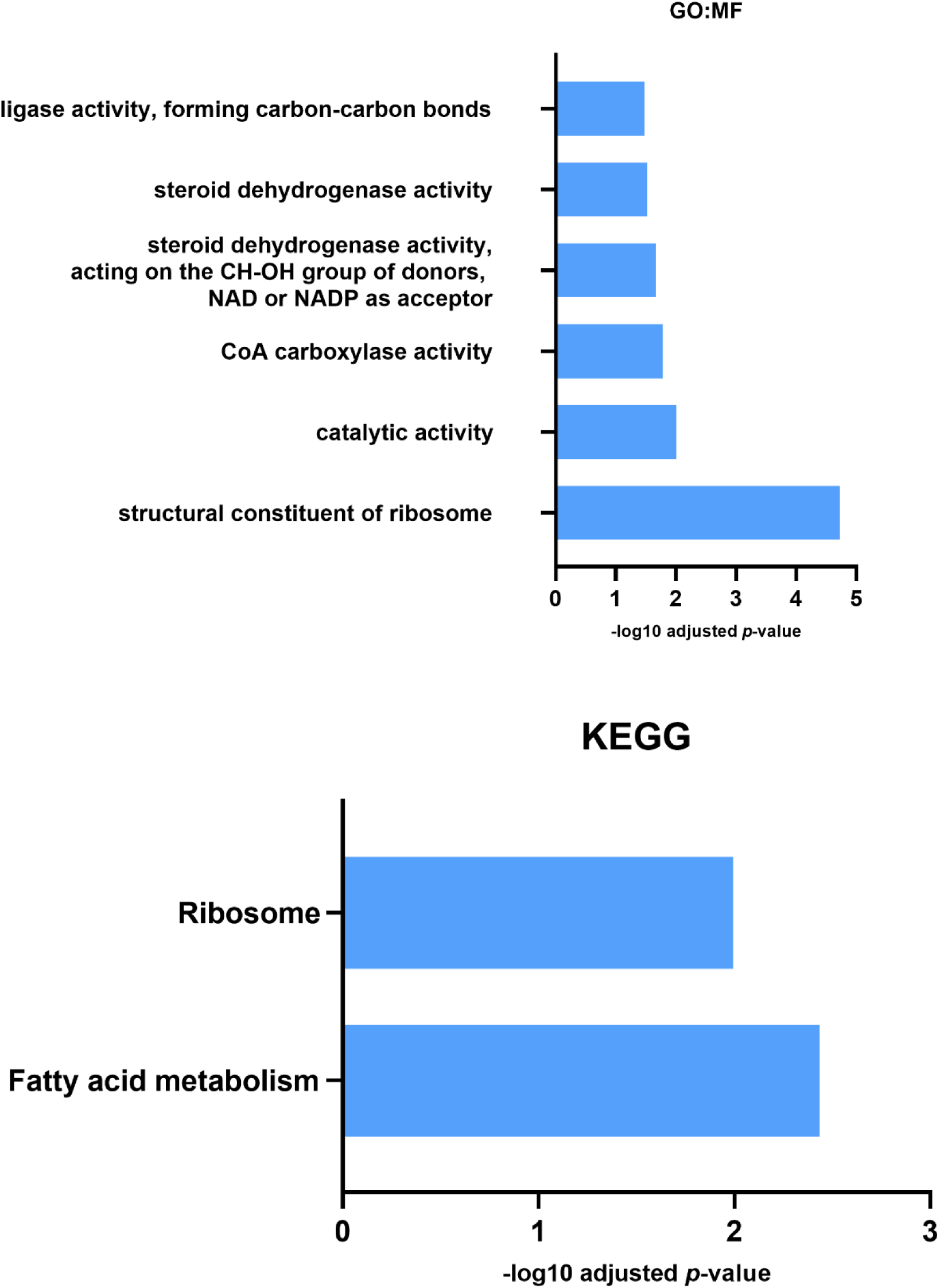
GO: Molecular Function and KEGG analysis for proteins reduced in expression in Pink1^-^ exercised flies compared with Pink1^-^ non-exercised flies. Both Molecular Function and KEGG enrichment analysis indicated a decrease in expression of fatty acid metabolic proteins in the exercised Pink1^-^ flies.

KEGG analysis identified fatty acid metabolism from proteins reduced in expression due to exercise, while GO:MF analysis highlighted CoA Carboxylase activity from the same protein data set. This reduction in the metabolism, and in particular the synthesis, of fatty acids can be contrasted with the KEGG analysis described earlier which identified elevated expression of fatty acid metabolism associated proteins in *Pink1^-^* flies compared with wild type flies. It can be interpreted that exercise reverses the change in *Pink1^-^* flies and returns the fatty acid metabolic profile back towards WT flies.

The two proteins upregulated with exercise in *Pink1^-^ D. melanogaster* were OCIA domain-containing protein 1 (OCIAD1) and dihydroorotate dehydrogenase (quinone) mitochondrial (DHODH), neither have previously been connected with exercise. OCIAD1 has been shown to localise to both endosomes and mitochondria and regulate pathways such as JAK/STAT, Notch and PI3K/AKT(Morel *et al*, 2013; Khadilkar *et al*, 2014; Li *et al*, 2020). OCIAD1 has been shown to regulate mitochondrial ETC activity via control of complex I activity, which showed an inverse association with OCIAD1 overexpression(Shetty *et al*, 2018). Deregulated OCIAD1 levels have been linked to mitochondrial dysfunction, interaction with BCL-2 and Alzheimer’s disease(Li *et al*, 2020).

DHODH is an inner mitochondrial membrane enzyme that catalyses the fourth step in *de novo* synthesis of pyrimidines(Cherwinski *et al*, 1995). A link between pyrimidine synthesis and mitochondrial morphology was shown with the addition of the drug leflunomide to muscle cells(Miret-Casals *et al*, 2018). The group showed that leflunomide inhibited DHODH by binding to its ubiquinone binding channel, thereby preventing the production of pyrimidine ribonucleotide uridine monophosphate (UMP). DHODH inhibition induced upregulation of mitofusions and subsequent mitochondrial elongation, by depleting the cellular pyrimidine pool. As ubiquinone is reduced to ubiquinol in the DHODH-mediated catalysis of dihydroorotate to orotate, and as ubiquinol is a substrate of respiratory complex III, DHODH is important for the ETS. DHODH deficiency has been reported to partially inhibit complex III and increase ROS generation(Fang *et al*, 2013).

The same *Pink1^-^ D. melanogaster* strain has previously been reported to have upregulated genes involved in nucleotide metabolism, which is also the case in brains of PD patients with *PINK1* mutations(Tufi *et al*, 2014). Genetic and pharmacological upregulation of nucleotide metabolism and scavenging pathways restored mitochondrial function caused by *PINK1* loss. Therefore, DHODH upregulation by exercise may act in a compensatory manner to manage metabolic stress due to the *Pink1^-^* phenotype.

### Exercise reduces the difference in levels of protein expression between *Pink1*^-^ and WT flies

Of the 516 proteins identified, 105 protein had different levels between *Pink1^-^* and WT non-exercised *D. melanogaster* (**Supp. Table 3**). Comparing the *Pink1^-^* exercised group to either WT non-exercised or exercised halved the number of differentially expressed proteins (55 and 56 proteins, respectively) (**Supp. table 2**). Qualitatively the heatmap of protein expression for *Pink1^-^* exercised *D. melanogaster* more closely resembles WT *D. melanogaster* (**Fig 4**). This suggests exercise can ameliorate the aberrant protein profile of *Pink1^-^ D. melanogaster* towards a more WT profile. In agreement, gene expression data from an exercised mouse model of arrhythmogenic cardiomyopathy, that originally showed a dysregulation of near 800 genes, showed partial restoration of gene expression with regular exercise, with greatest remedial effects on proteins involved in inflammation and oxidative phosphorylation(Cheedipudi *et al*, 2020).

**Figure 4.**
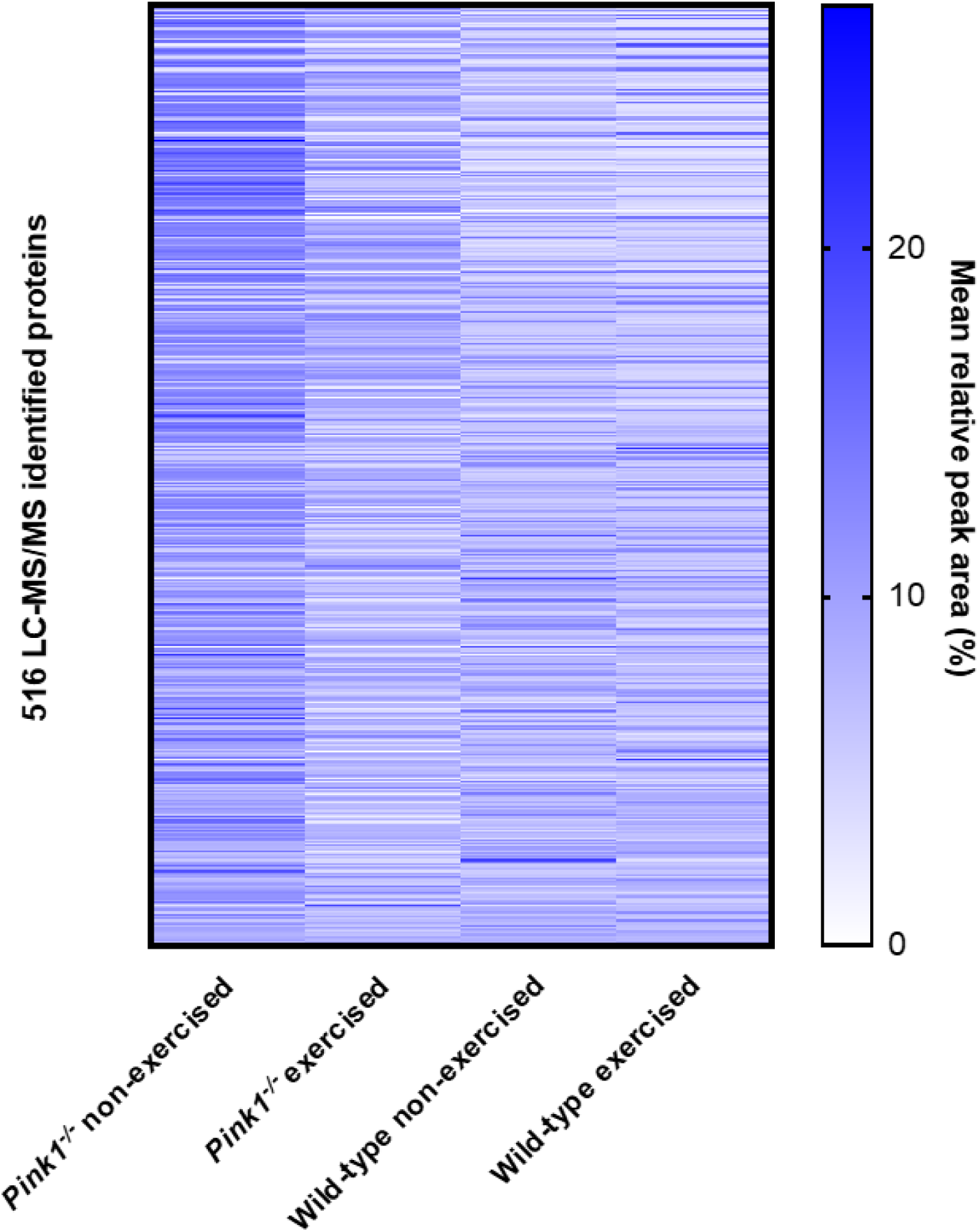
Heat map of protein expression levels (for proteins identified among all groups), determined by label-free mass spectrometry of mitochondrial fractions. Qualitatively, the identified Pink1^-^ exercised fly proteome more closely resembles the two WT fly proteomes than does the Pink1^-^ non-exercised fly proteome.

### Network analyses of differentially expressed proteins

STRING database network analysis complemented the results seen with gProfiler. The protein-protein interaction (PPI) networks were generated using STRING, and (**Fig 5A**) shows that *Pink1^-^* exercised compared with the WT exercised flies have a differentially expressed proteins (DEPs) network that consisted of 49 nodes and 200 edges with average node degree 8.16 and PPI enrichment p-value of (P < 1.0e-16).

**Figure 5.**
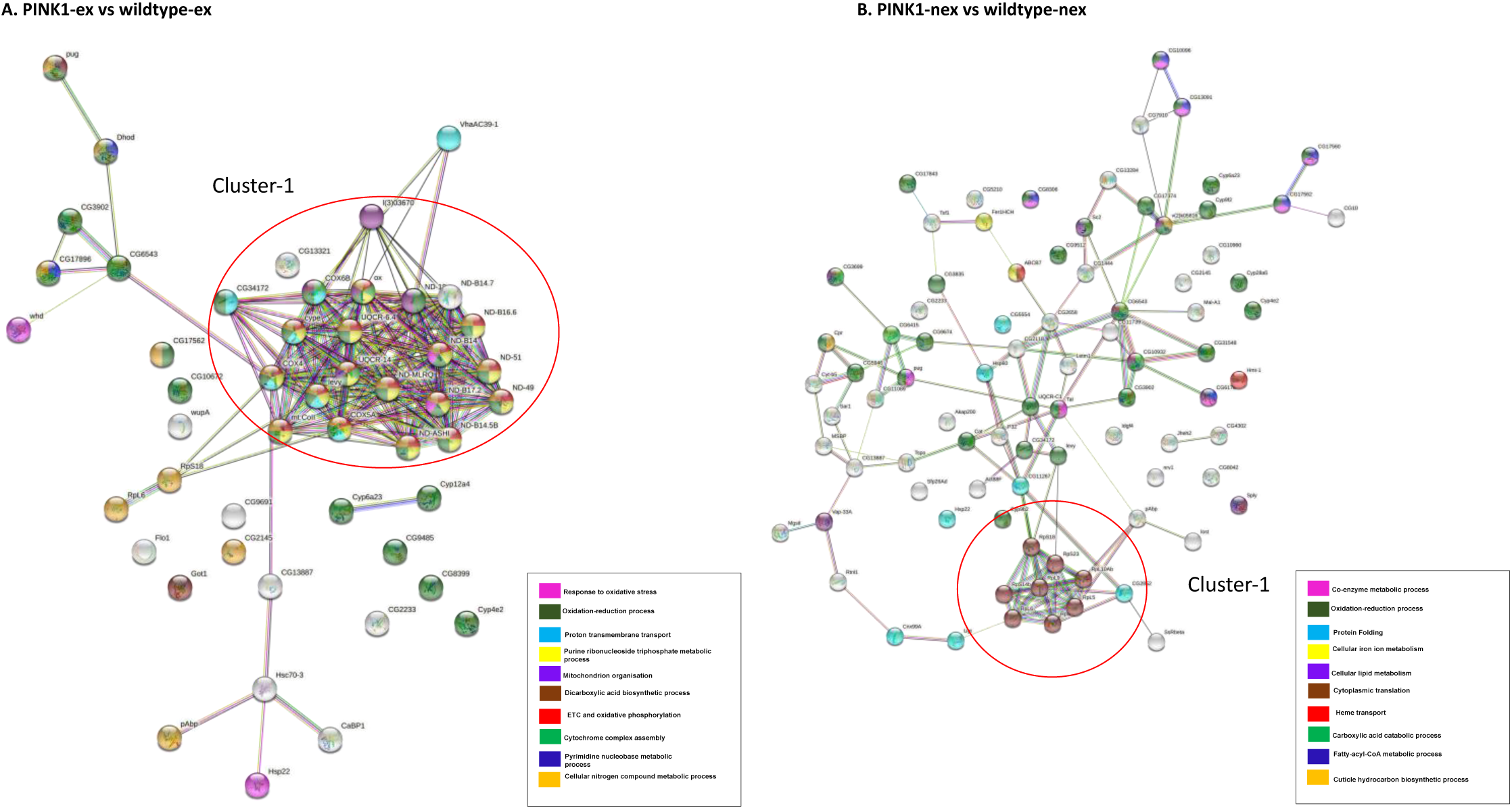
Network Analysis of differentially expressed proteins. A) Pink1^-^ exercised vs WT exercised; B) Pink1^-^ non-exercised vs WT non-exercised. Networks analysed using STRINGdb. The nodes are coloured according to the processes (legend) that the proteins are involved in by using GO Terms for Biological Processes. The edge shows type of interactions, experimentally determined interactions are pink and those obtained from databases are sky blue. Predicted interactions such as gene neighbourhood are blue, green and red for gene co-occurrence, gene neighbourhood and gene fusions. Co-expression interactions are shown in black, text-mining interactions are shown in light green and protein homology edges are purple.

Most of the proteins in the network have downregulated expression, with the following proteins found to be upregulated: endoplasmic reticulum chaperone BiP, enoyl-CoA hydratase short chain, glutamine synthetase, protein disulphide isomerase, methylmalonate-semialdehyde dehydrogenase, polyadenylate-binding protein, poly(U)-specific endoribonuclease, glutamine synthetase, flotillin-1, heat shock protein 22, phosphotidate cytidylyltransferase, fatty acyl-CoA reductase and dihydroorotate dehydrogenase.

The DEPs were found to be part of the ETC and OXPHOS processes, transmembrane transport and oxidative-reduction processes. The highly connected network consisted of proteins from complex I: ND-18, ND-MLRQ, ND-ASHI, ND-B14.5B, ND-B14, ND-B16.6, ND-49, ND-51, ND-B17.2, ND-B14.7; complex IV: COX6B, COX4, mt: COII, COX5A; complex III: OX, and other proteins CYPE, UQCR-6.4, UQCR-14, levy and 40S ribosomal proteins S18. This cluster of proteins are downregulated in the PINK1^-^ exercised. (**Fig 5B**) shows the *Pink1^-^* non-exercised compared with the WT non-exercised PPI network consisted of 86 nodes and 121 edges with average node degree 2.81 and PPI enrichment p-value (P < 1.0e-16). In contrast to *Pink1^-^* exercised versus WT exercised all DEPs were upregulated, except for cytochrome c oxidase subunit 4, cyclope isoform A, flightin isoform B, cytochrome b-c1 complex subunit 7, NADH dehydrogenase 1 alpha subcomplex 12, Troponin 1, NADH dehydrogenase 18, cytochrome c oxidase subunit, cytochrome P450, NADH dehydrogenase 1 beta subcomplex subunit 8, cytochrome c oxidase subunit 5A, NADH dehydrogenase B14 and levy isoform A.

The DEPs were found to be a part of oxidation-reduction processes, fatty-acyl-CoA metabolic processes, translation and protein folding. The identification of DEPs from fatty-acyl-CoA metabolic process is concurrent with the enrichment analysis presented in (**Fig. 3**). The highly connecting nodes in this network is that of ribosomal proteins, RPS18, RPS23, RPS23, RPS7, RPS14b, RPL6, RPL5, RPL10AB and RPL9 involved in translation that are upregulated.

Similarly, for (**Fig 6A-C**), three additional PPI networks were generated for DEPs in pairwise group comparisons. For *Pink1^-^* exercised flies compared with *Pink1^-^* non-exercised flies the PPI network consisted of 43 nodes and 29 edges with average node degree 1.35 and PPI enrichment p-value <1.62e-06 (**Fig 6A**). For *Pink1^-^* non-exercised flies compared with WT exercised flies the network has 107 nodes and 341 edges with average node degree 6.37 and PPI enrichment p-value (P <1.0e-16) (**Fig 6B**). For the final PPI network, *Pink1^-^* exercised flies compared with WT non-exercised flies, there are 44 nodes and 40 edges with average node degree 1.82 and PPI enrichment p-value 6.79e-11 (**Fig 6C**). For each of the three PPI networks, the number of edges is larger than expected and the nodes were more connected than for a random PPI network of the same size.

**Figure 6.**
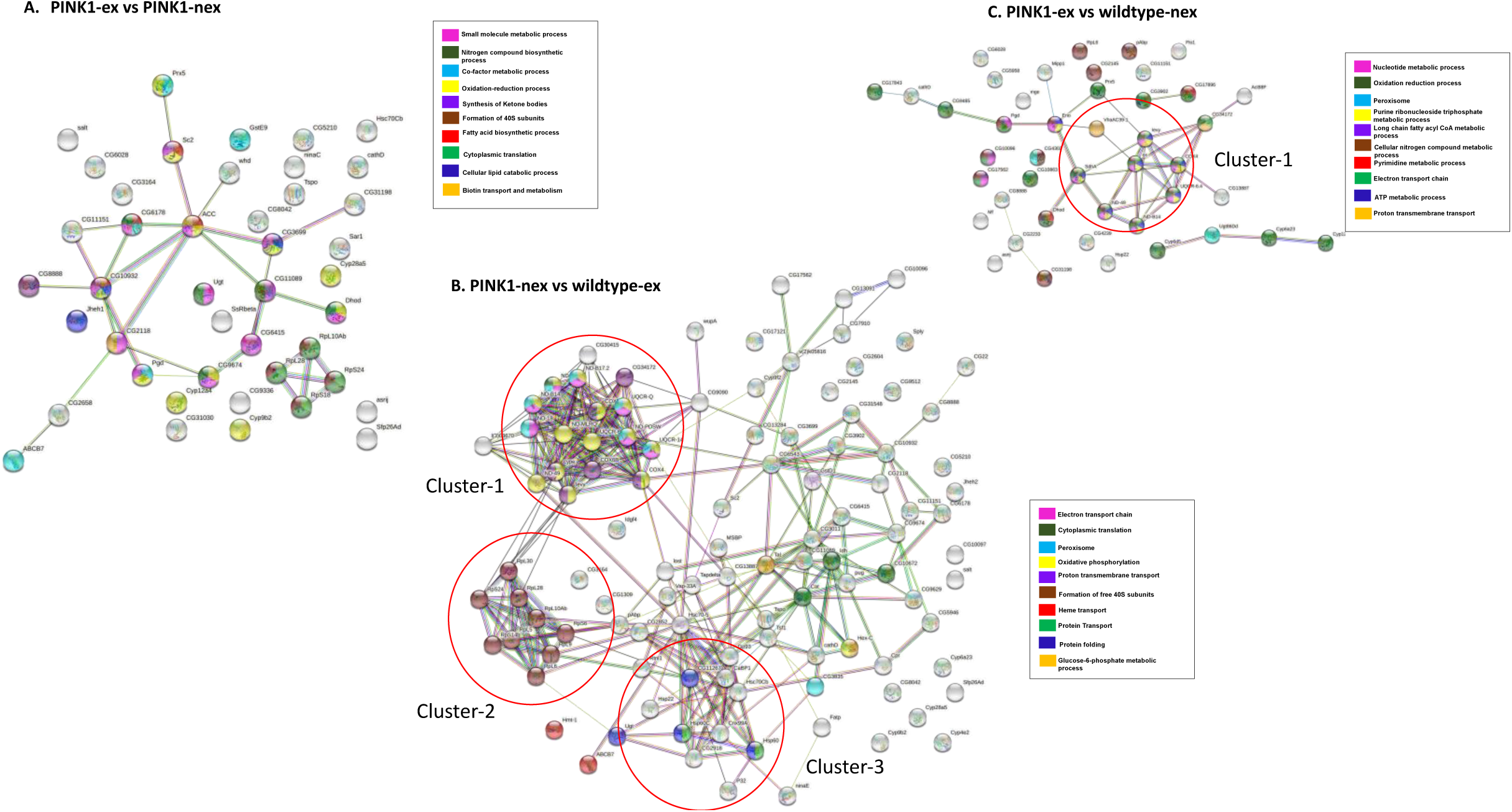
Network Analysis of differentially expressed proteins. A) Pink1^-^ exercised vs Pink1^-^ non-exercised; B) Pink1^-^ non-exercised vs WT exercised; C) Pink1^-^ exercised vs WT non-exercised. Networks analysed using STRINGdb. The nodes are coloured according to the processes (legend) that the proteins are involved in by using GO Terms for Biological Processes. The edge shows type of interactions, experimentally determined interactions are pink and the one obtained from databases are sky blue. Predicted interactions such as gene neighbourhood are blue, green and red for gene co-occurrence, gene neighbourhood and gene fusions. Co-expression interactions are shown in black, text-mining interactions are shown in light green and protein homology edges are purple.

In *Pink1^-^* exercised flies versus *Pink1^-^* non-exercised flies all DEPs were downregulated except for OCIA domain containing protein 1 and dihydroorotate dehydrogenase (**Fig 6A**). The *Pink1^-^* exercised flies and *Pink1^-^* non-exercised flies DEPs were found to be involved in formation of 40S ribosomal subunit, oxidation reduction processes, synthesis of ketone bodies, translation and cellular lipid catabolic processes. *Pink1^-^* non-exercised flies and WT exercised flies DEPs were found to be involved in ETC, translation, peroxisome, formation of 40S subunits and protein folding (**Fig 6B**). The *Pink1^-^* exercised versus WT non-exercised were involved in purine ribonucleotide triphosphate metabolic processes, ETC and peroxisomes (**Fig 6C**).

PINK1^-^ non-exercised compared with wildtype exercised showed three highly connected clusters (**Fig 6B**). Cluster 1 had proteins from complex I and complex IV proteins that are downregulated like that of *Pink1^-^* exercised compared with the WT exercised flies and Cluster 2 has ribosomal proteins that are upregulated, similar to that of *Pink1^-^* non-exercised compared with the WT non-exercised PPI network. Cluster 3 has proteins UGT, HSP22, HSP60C, CABP1, GP93, HSC70-5, HSC70Cb, HSP60A, RTNL1 and CNX99A, proteins involved in protein control and folding which are found to be upregulated.

## Conclusion

A picture of global proteomic changes made in response to exercise was obtained. By using two methodologies to assess the mitochondrial proteome we were able to measure changes in an organelle whose function in exercise, and dysfunction in Parkinson’s disease, is crucial.

2D-GE MS analysis used to compare WT exercised flies and control flies identified five out of seven proteins with altered expression, the majority of which have roles in metabolism and bioenergetics. 2D-GE MS data comparison between *Pink1^-^* exercised and *Pink1^-^* non-exercised flies revealed several proteins with decreased levels of expression in response to exercise.

GO and KEGG analyses validated our mitochondrial isolation methodology by identifying the enrichment of mitochondrial processes and pathways. In the four comparisons that we made; proteins involved in bioenergetics had changed expression. GO, KEGG and STRING network analysis of these proteome comparisons identified enrichment of bioenergetic pathways. The most significant pathways included oxidation-reduction, fatty acid metabolism, and folate metabolism which are associated with PD. Our data point to exercise aiding normalisation of these pathways. Specific proteins in the pathways may be candidates to develop therapeutic approaches in PD.

## Author contributions

BE performed data analysis, assisted with experiments, and wrote the manuscript, TLI performed experimental work and helped prepare the manuscript, GK performed STRING analyses and helped prepare the manuscript, JRI developed and constructed the ICE machine for fly exercise, NM generated the flies used for these experiments and was consulted on all aspects of the fly work, LC directed the research, supervised experiments, provided reagents and prepared the manuscript.

The authors declare they have no conflict of interest.

## Appendix

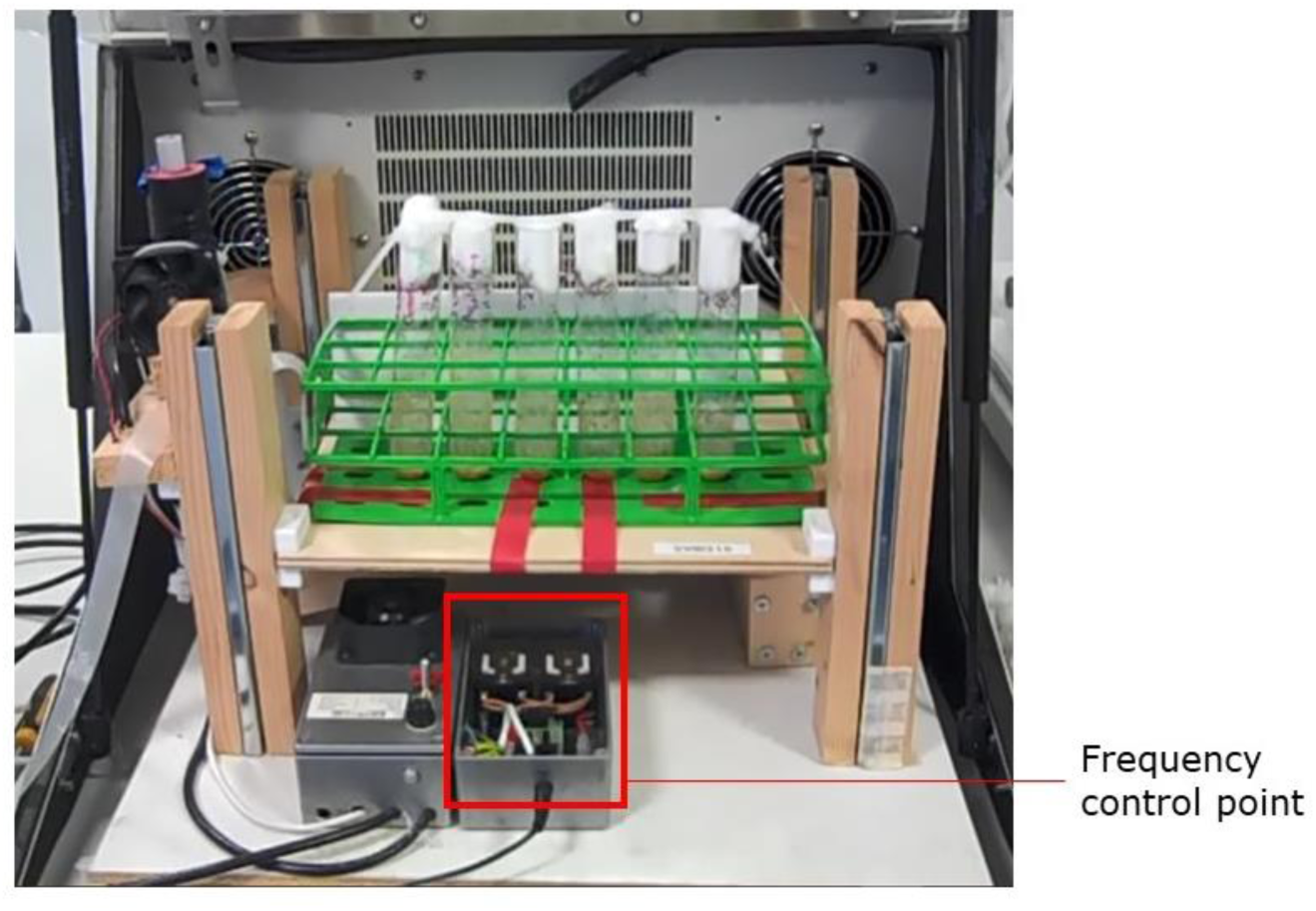

### Ingram counter-balanced exerciser

Our bespoke Ingram Counter-balanced Exercise (ICE) machine was adapted from the PT design of Piazza *et al.,* by Mr. John Ingram (TI’s father)(Piazza *et al*, 2009). The exerciser fits into an incubator with internal dimensions of 48 x 48 x 35cm and provides a vertically moving tray measuring 30 x 34cm. A solenoid-powered counterbalanced lever causes the tray to be lifted 3cm. The tray is lifted and immediately dropped every 15 seconds.

Counterbalancing the lever are a series of springs, which can be adjusted to allow lifts of up to 3.0kg. Springs efficiently store and release energy enabling a more rapid drop than would be the case if weights were used. They also reduce the size and overall weight of the device.

Lift is provided by a solenoid wound onto a nylon core and fixed to the body of the exerciser. The steel armature passing through the solenoid is attached to the counterbalanced lever. Activation of the solenoid causes the armature to rise which lifts the lever.

Solenoid activation is controlled by an astable timer. This triggers a relay to pulse the applied AC voltage. Two capacitors acting as a loss-less resistor allow the voltage to be reduced without producing excess heat, before it is rectified, smoothed and finally applied to the solenoid. Powering with DC current causes less vibration and heat generation in the solenoid but, as the capacitors and the solenoids both work more efficiently at lower temperatures, any excess heat is subsequently dissipated by proximate fans. Using a pulse of current as described rather than the discharge from a large capacitor to activate the solenoid saves space, it is also safer as there is far less stored energy.

## *Pink1* manuscript supplemental data

**Supplemental table 1.**
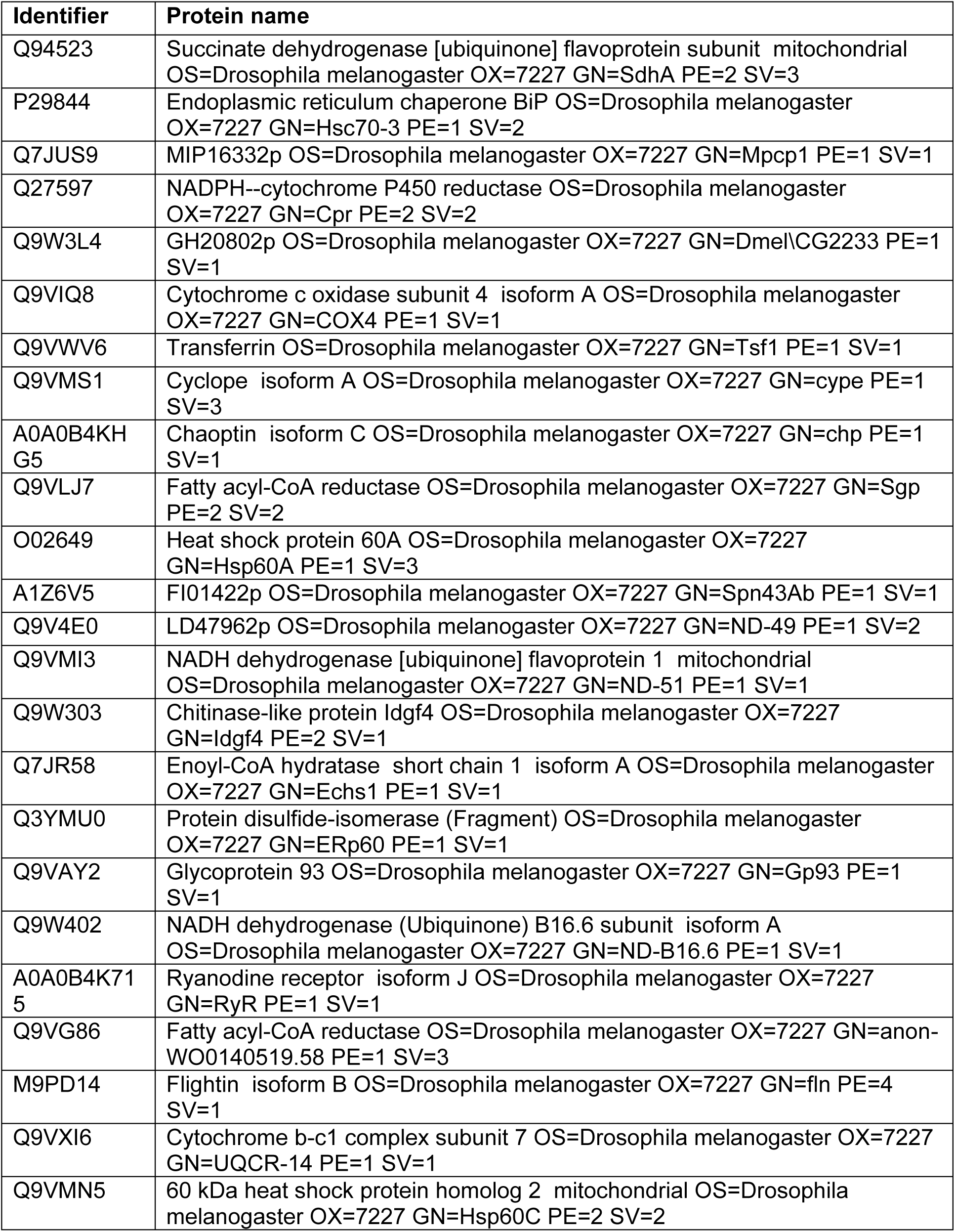
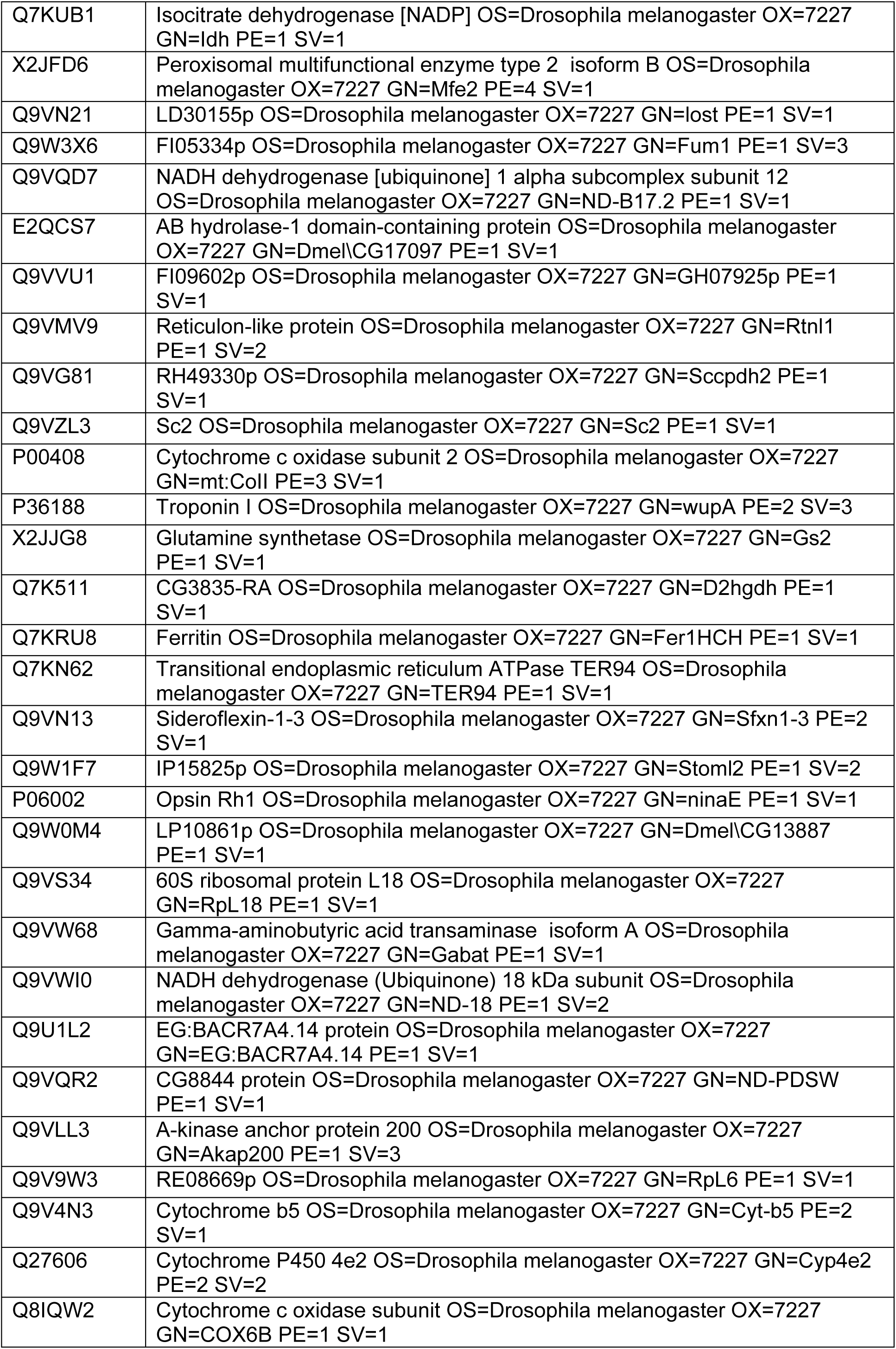
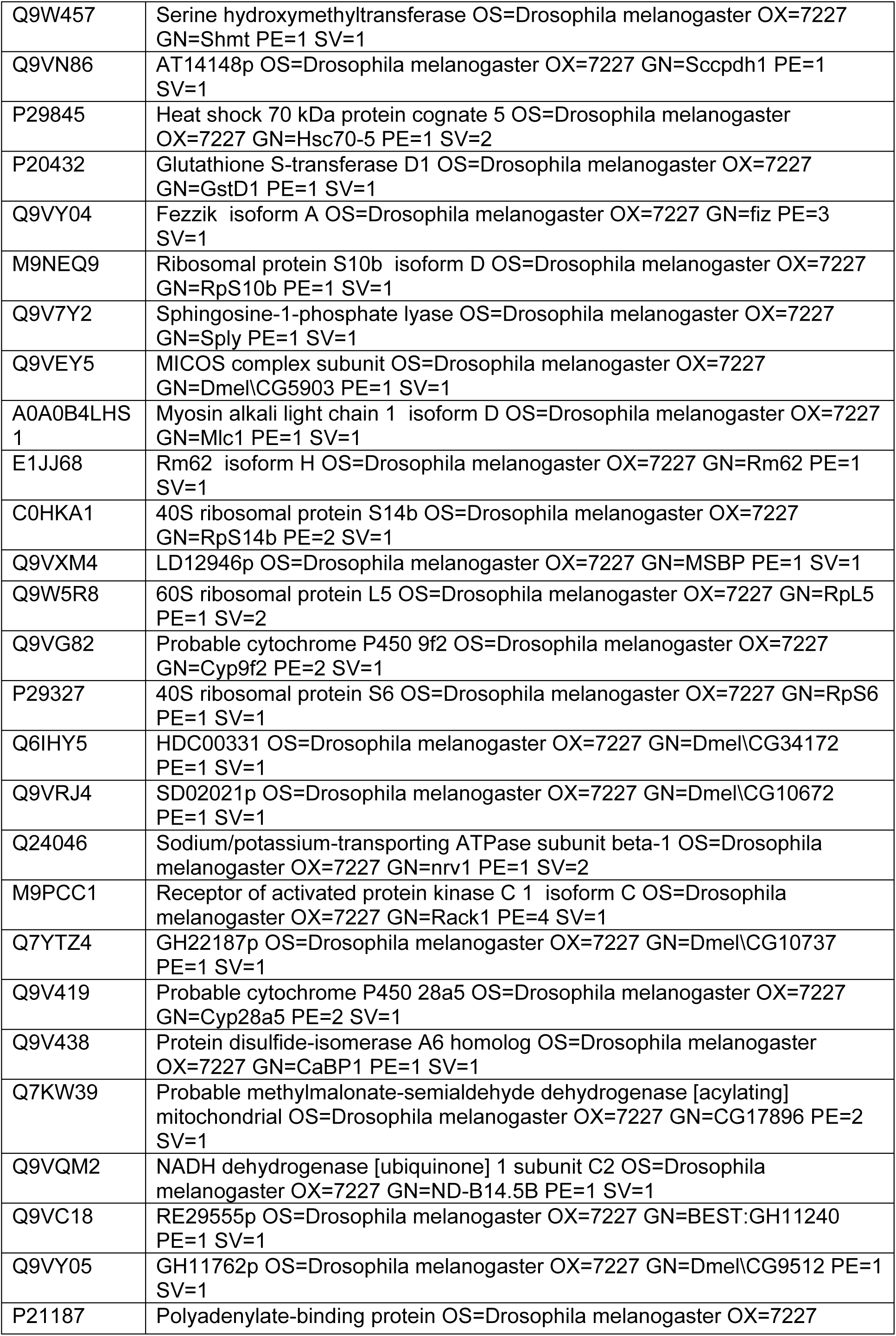
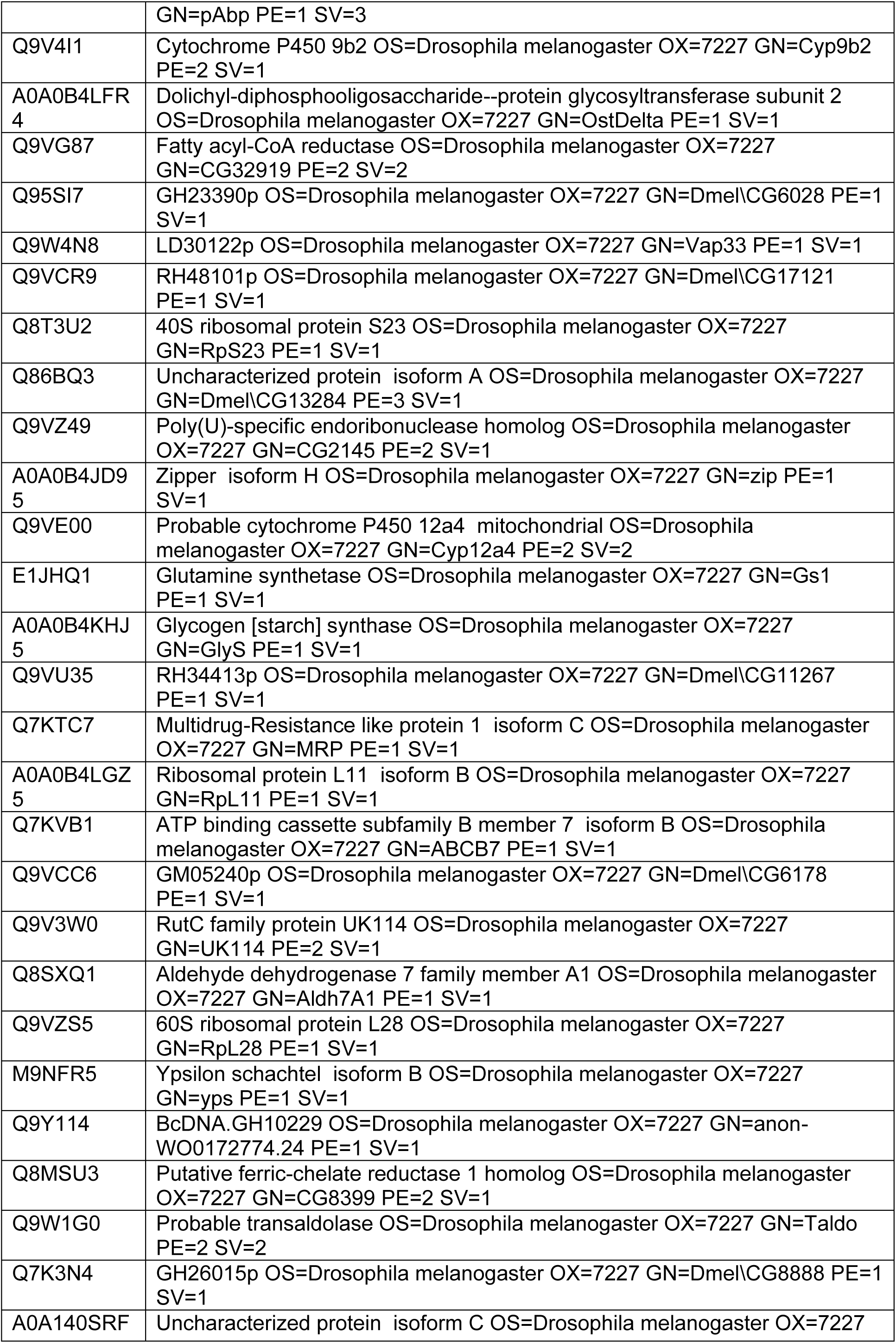
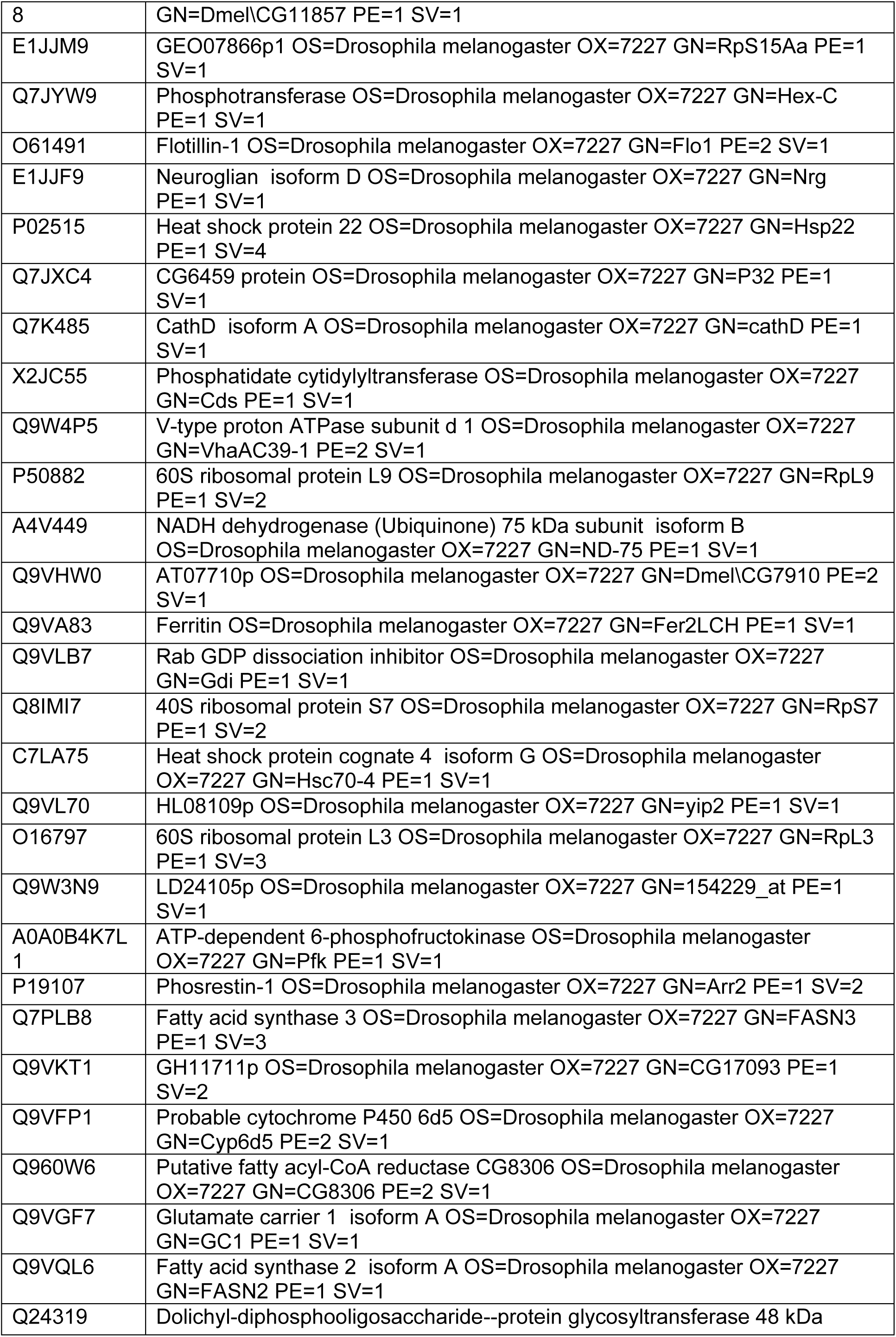
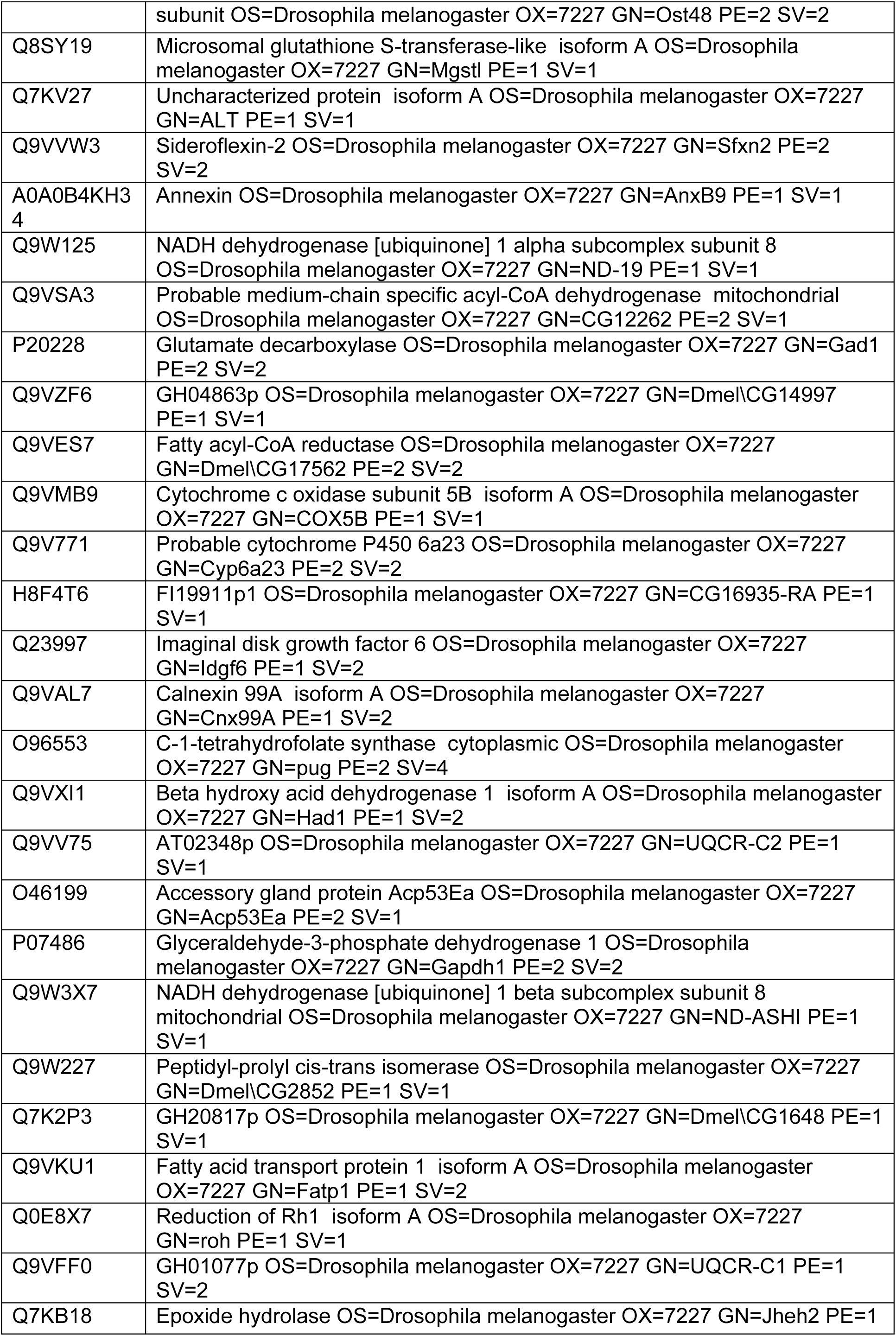
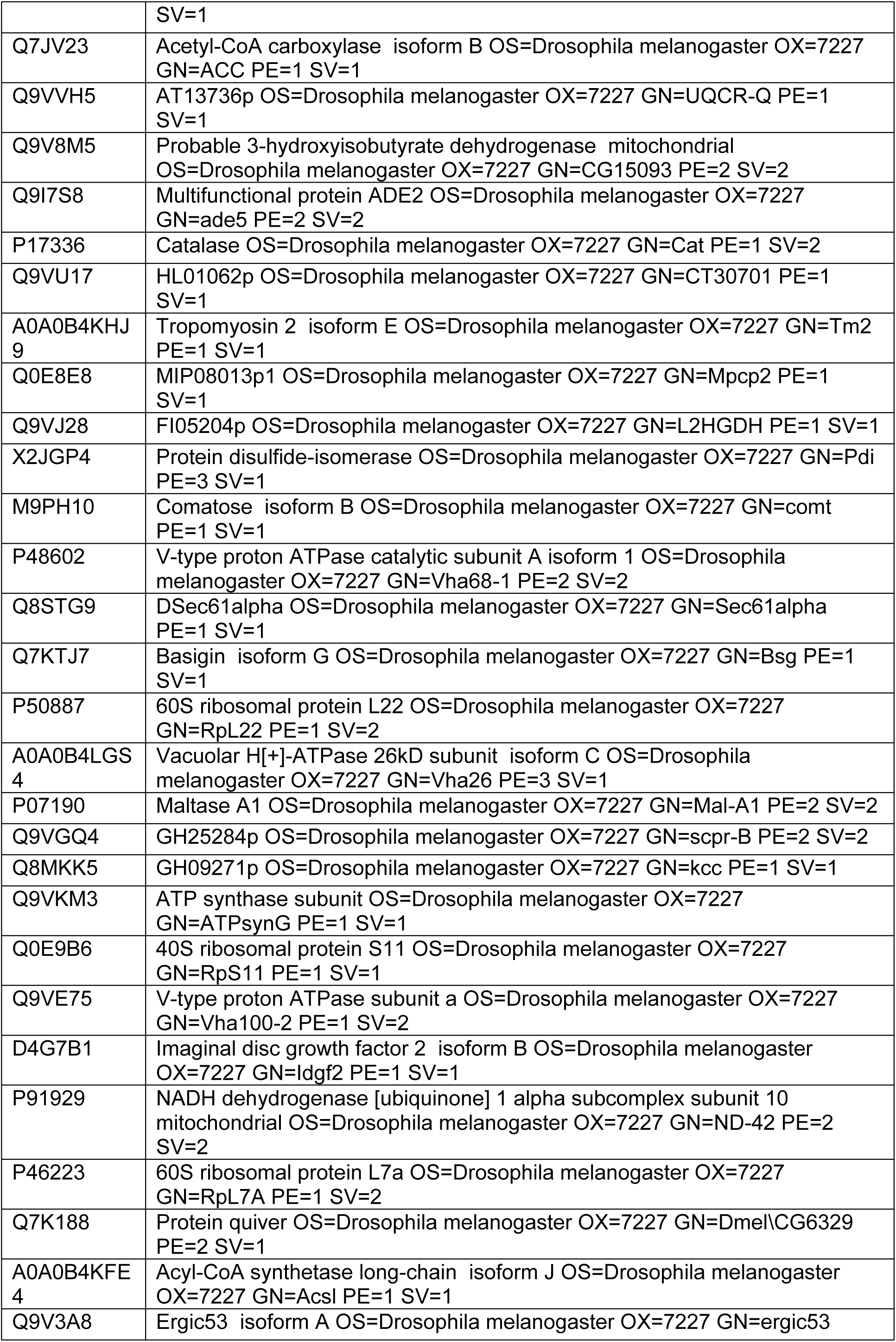
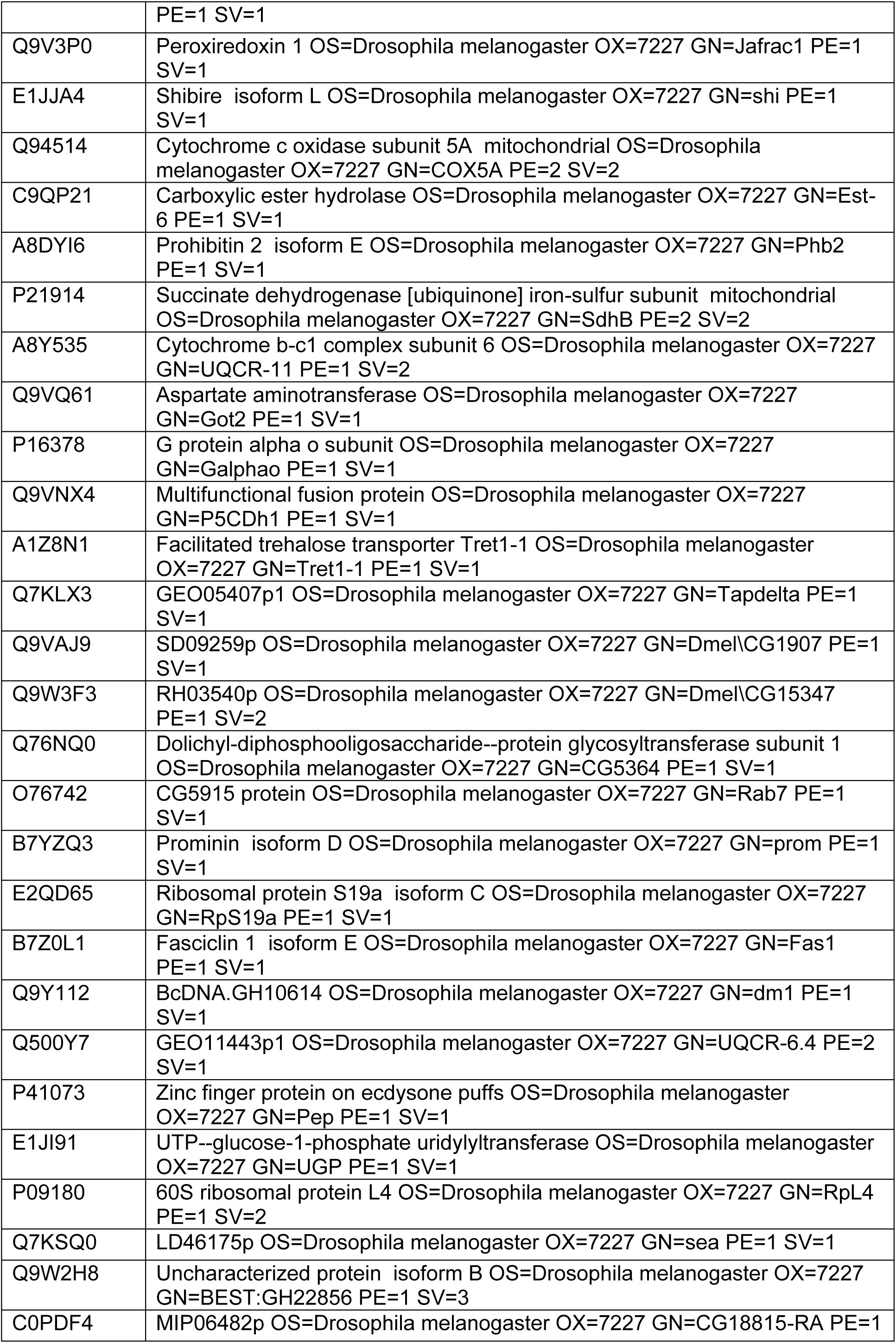
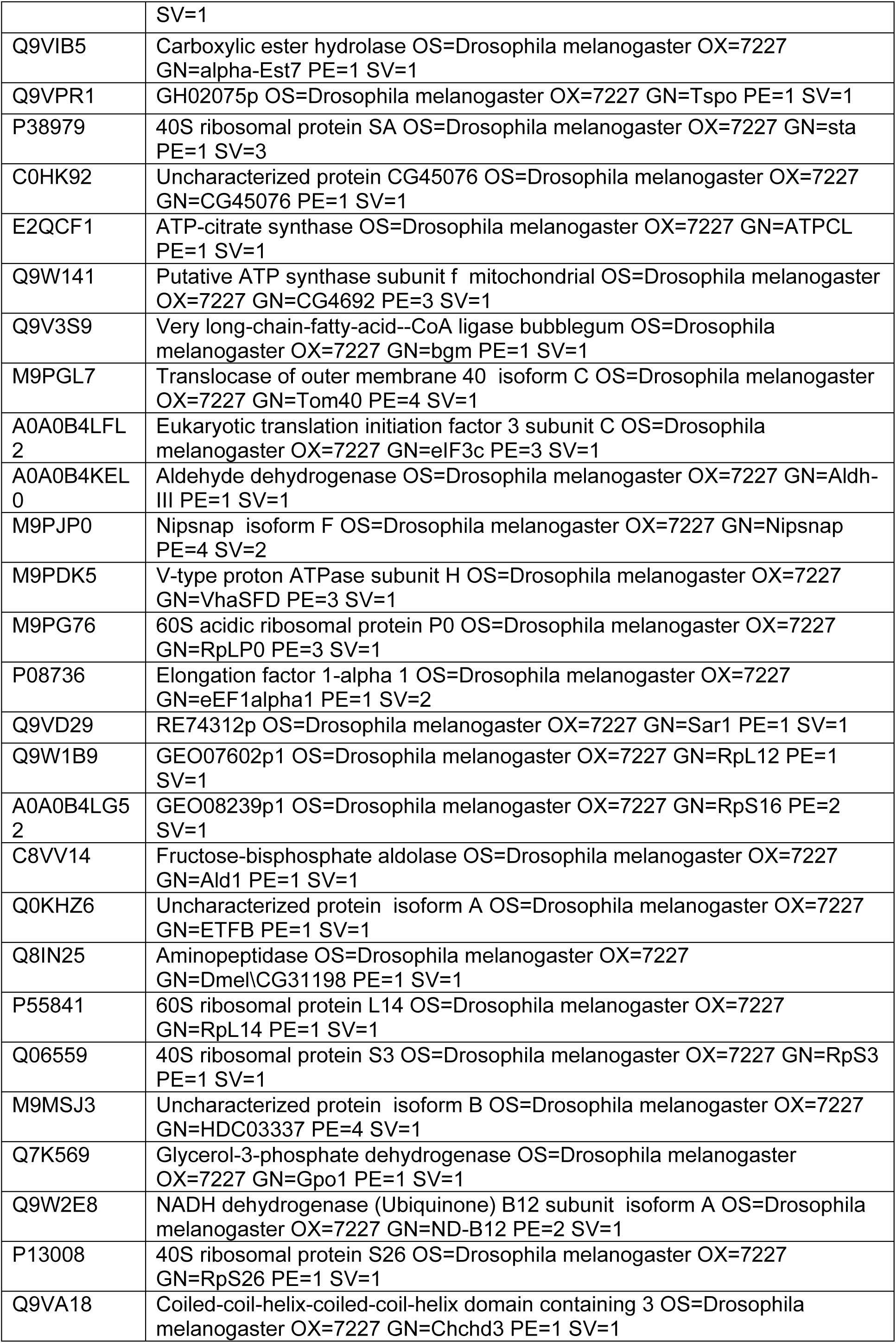
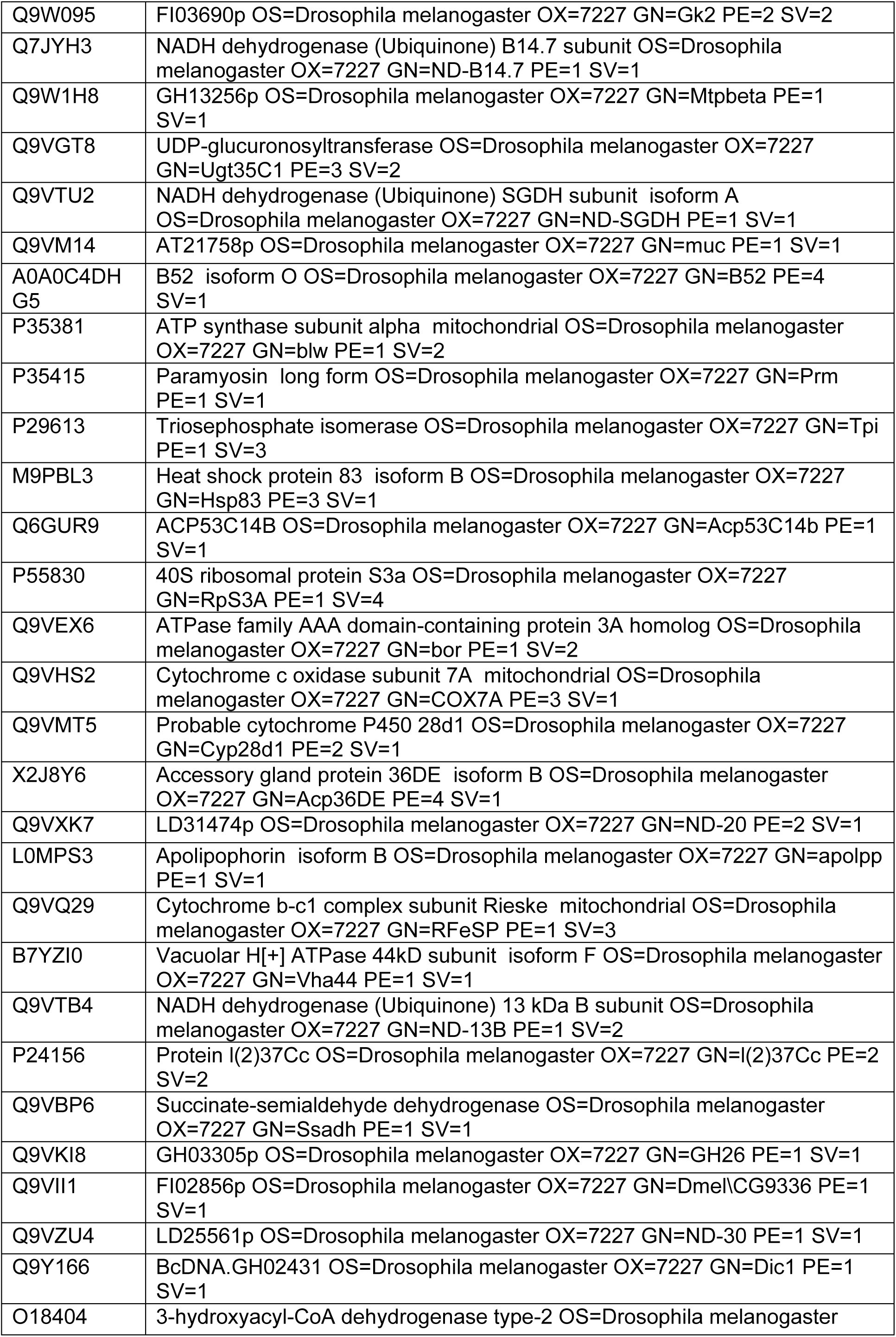
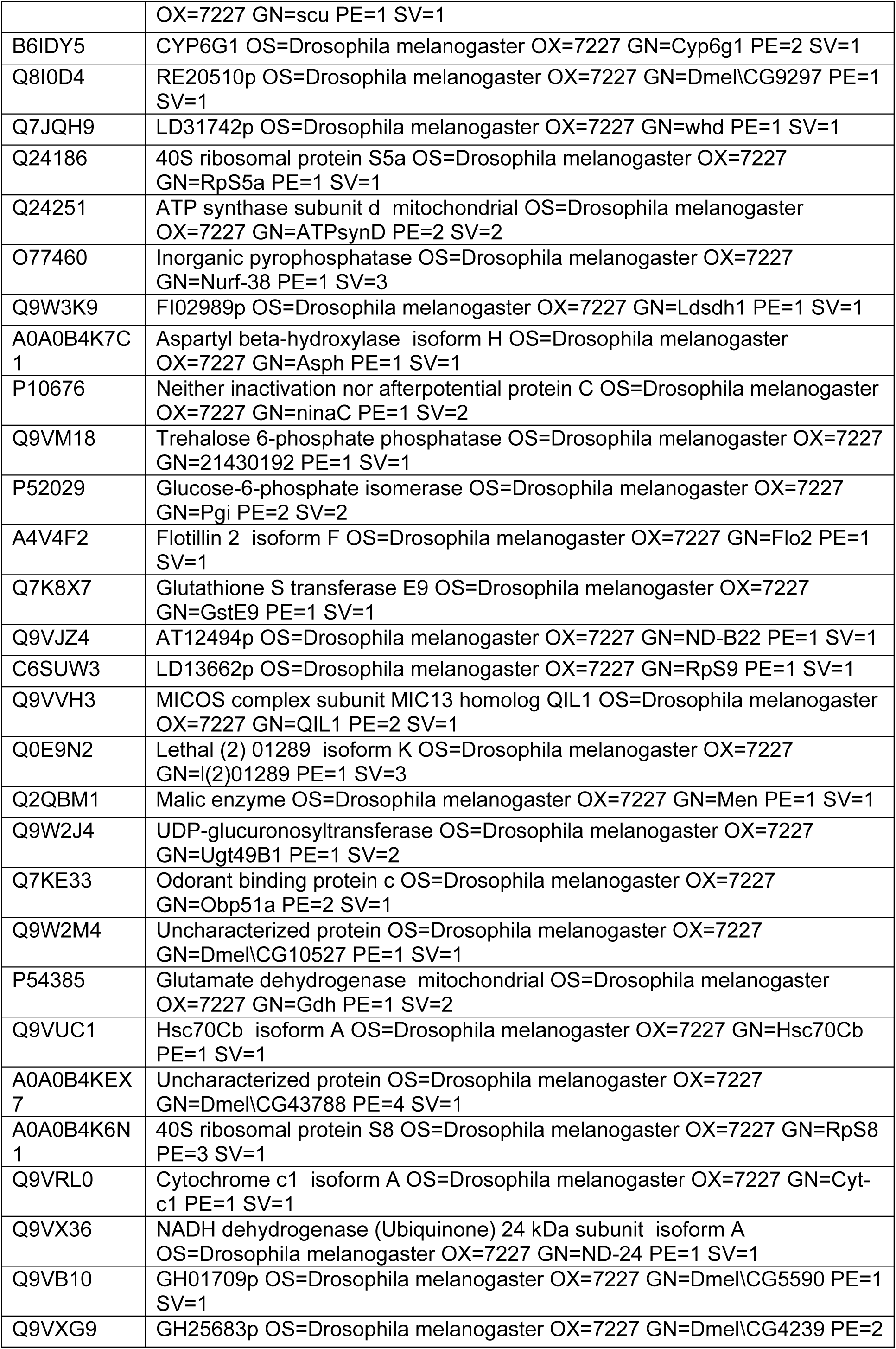
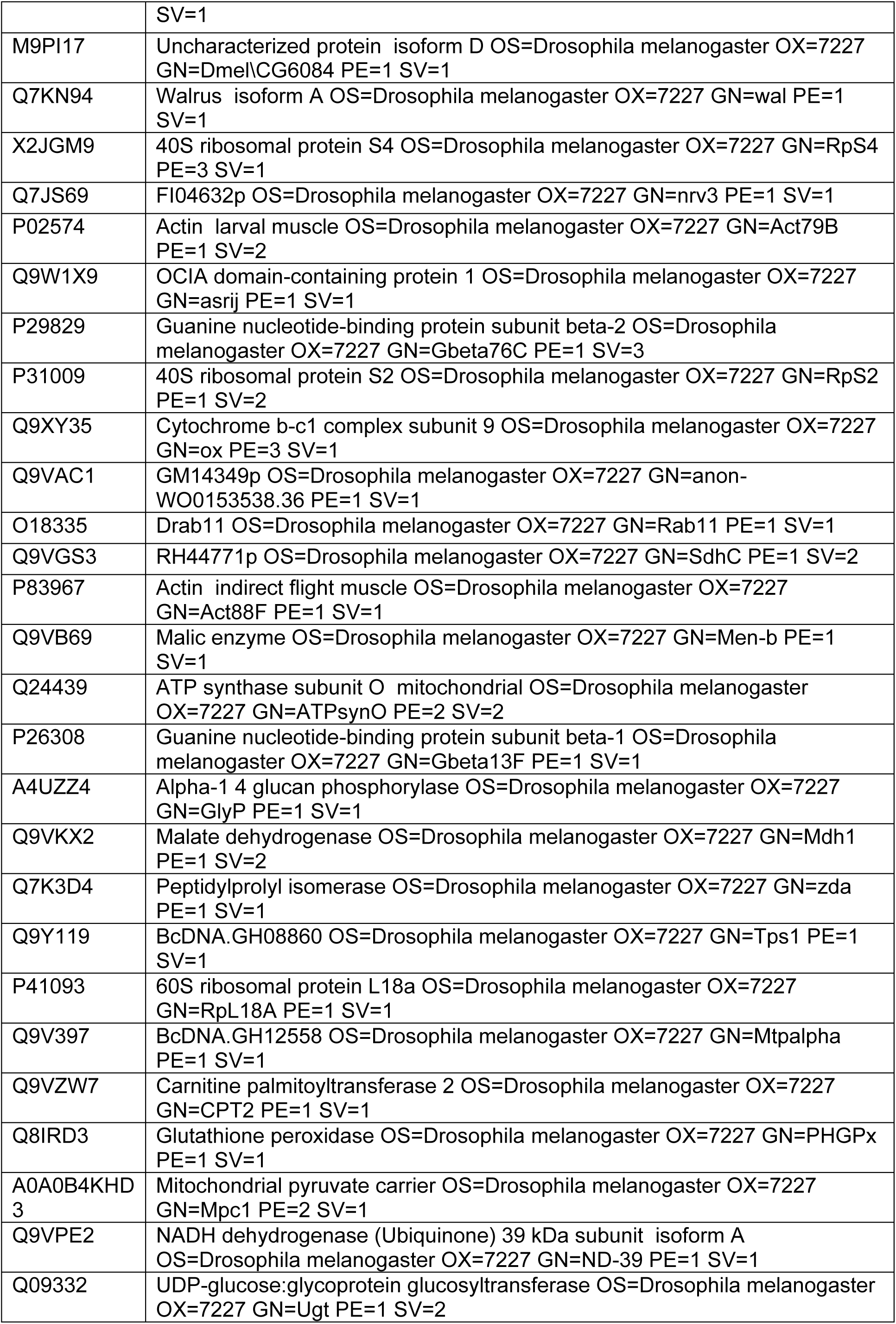
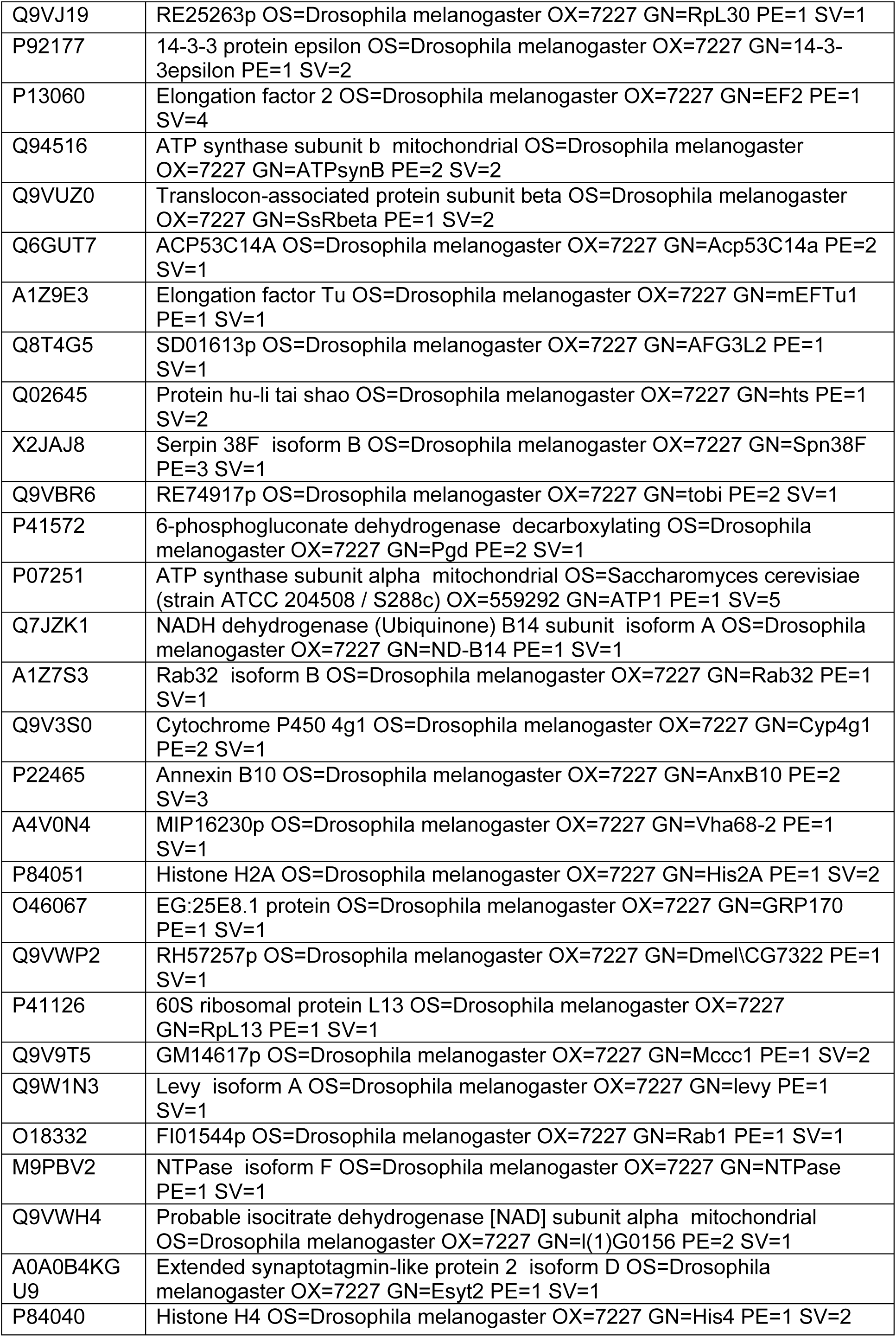
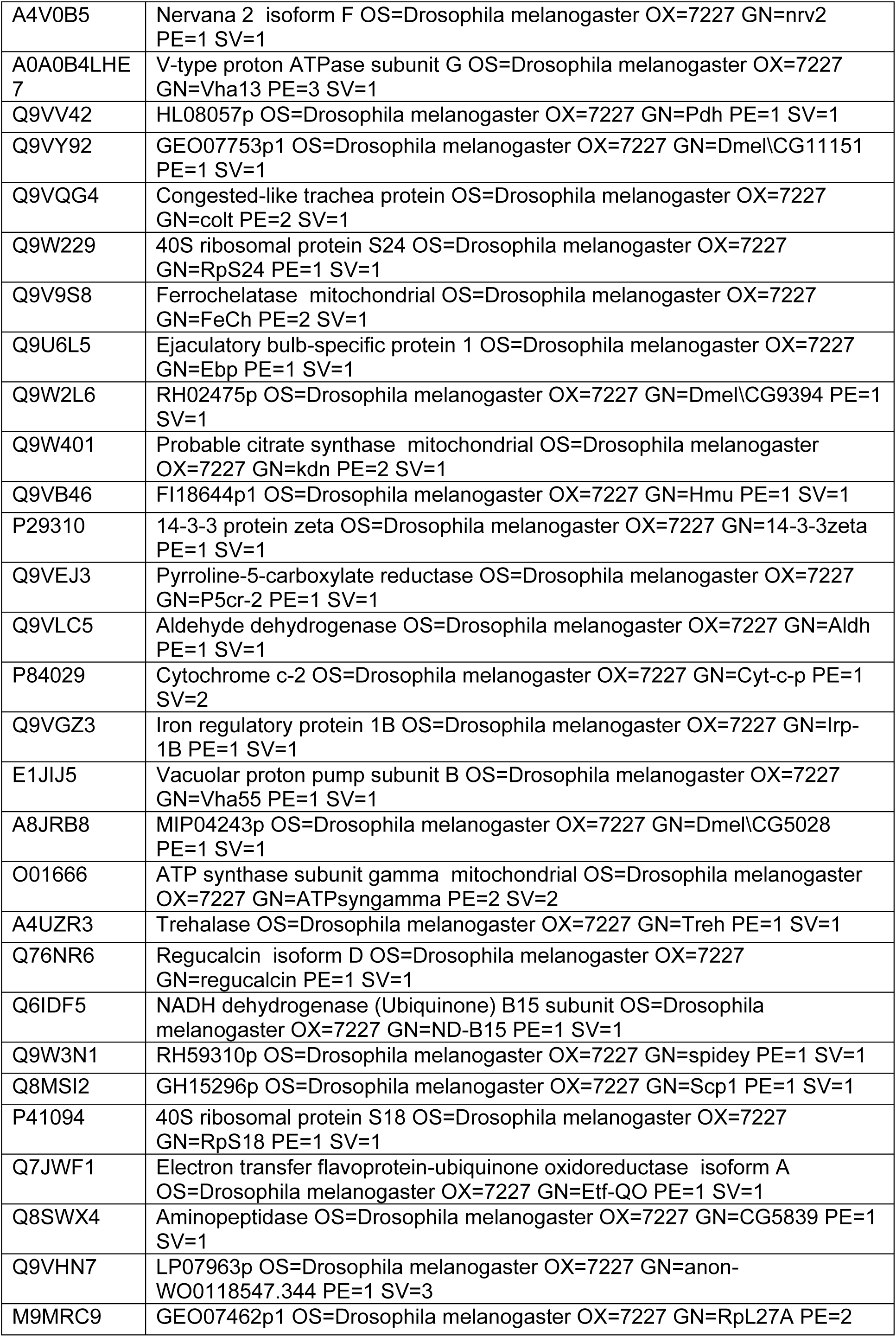
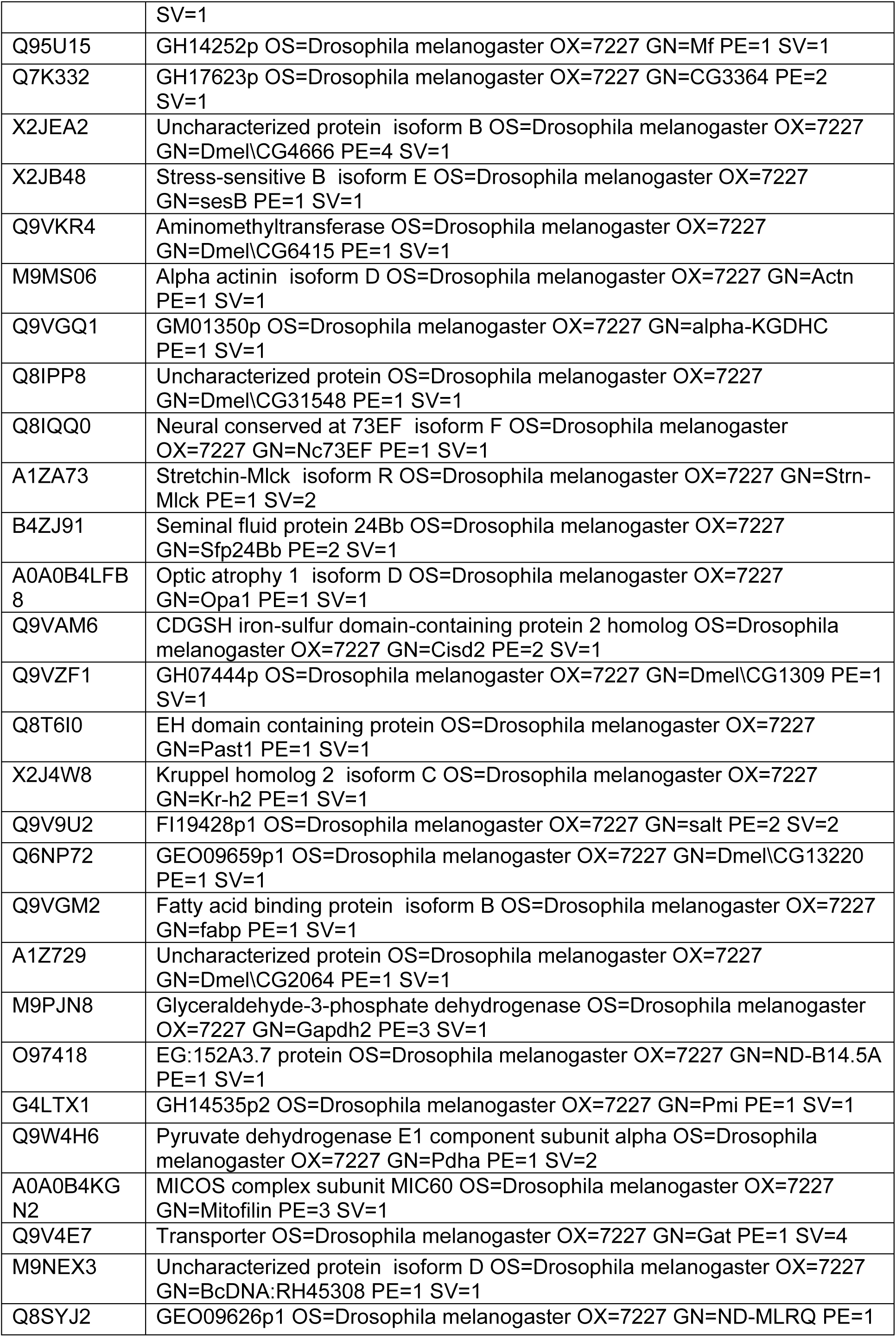
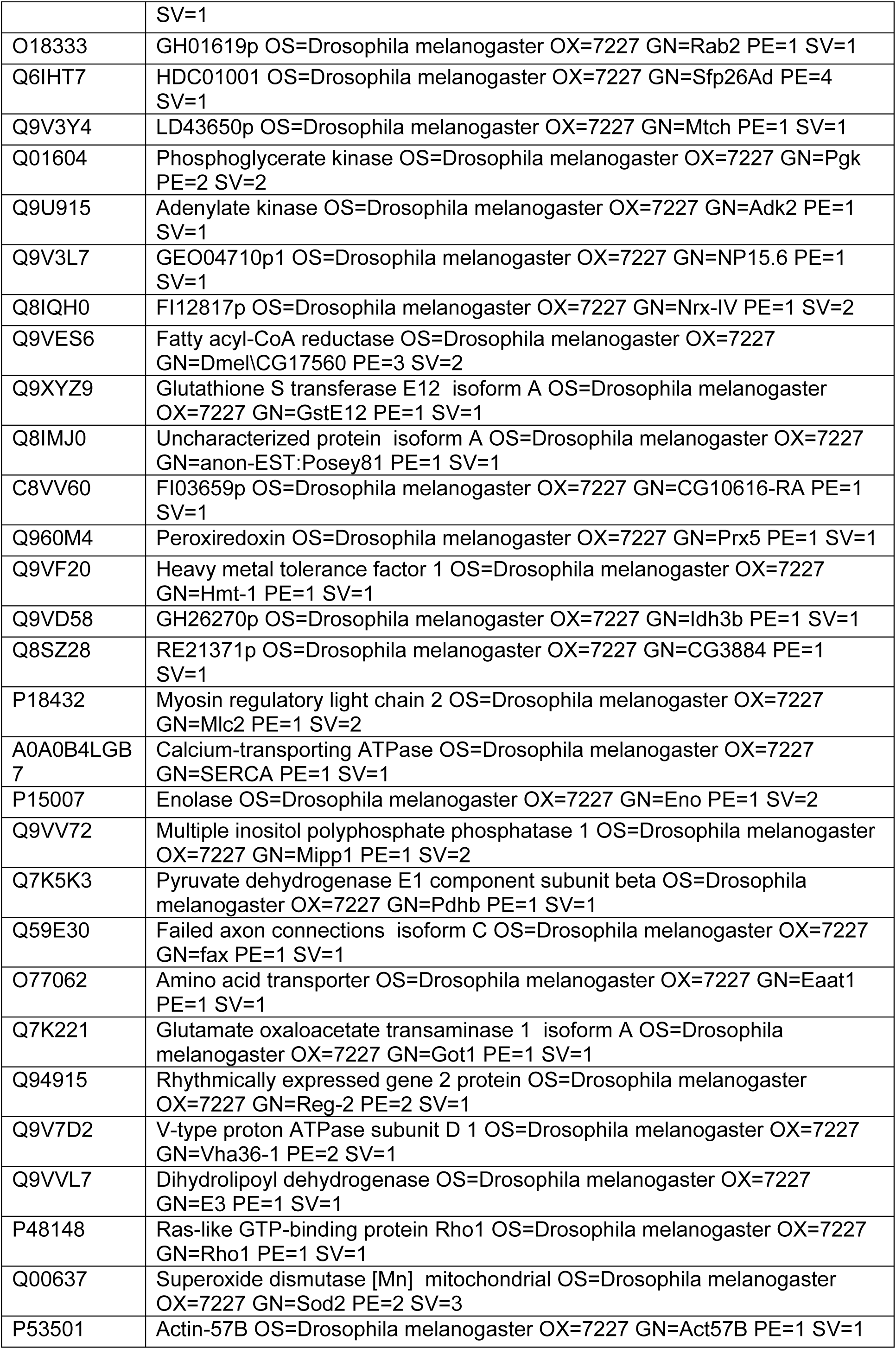
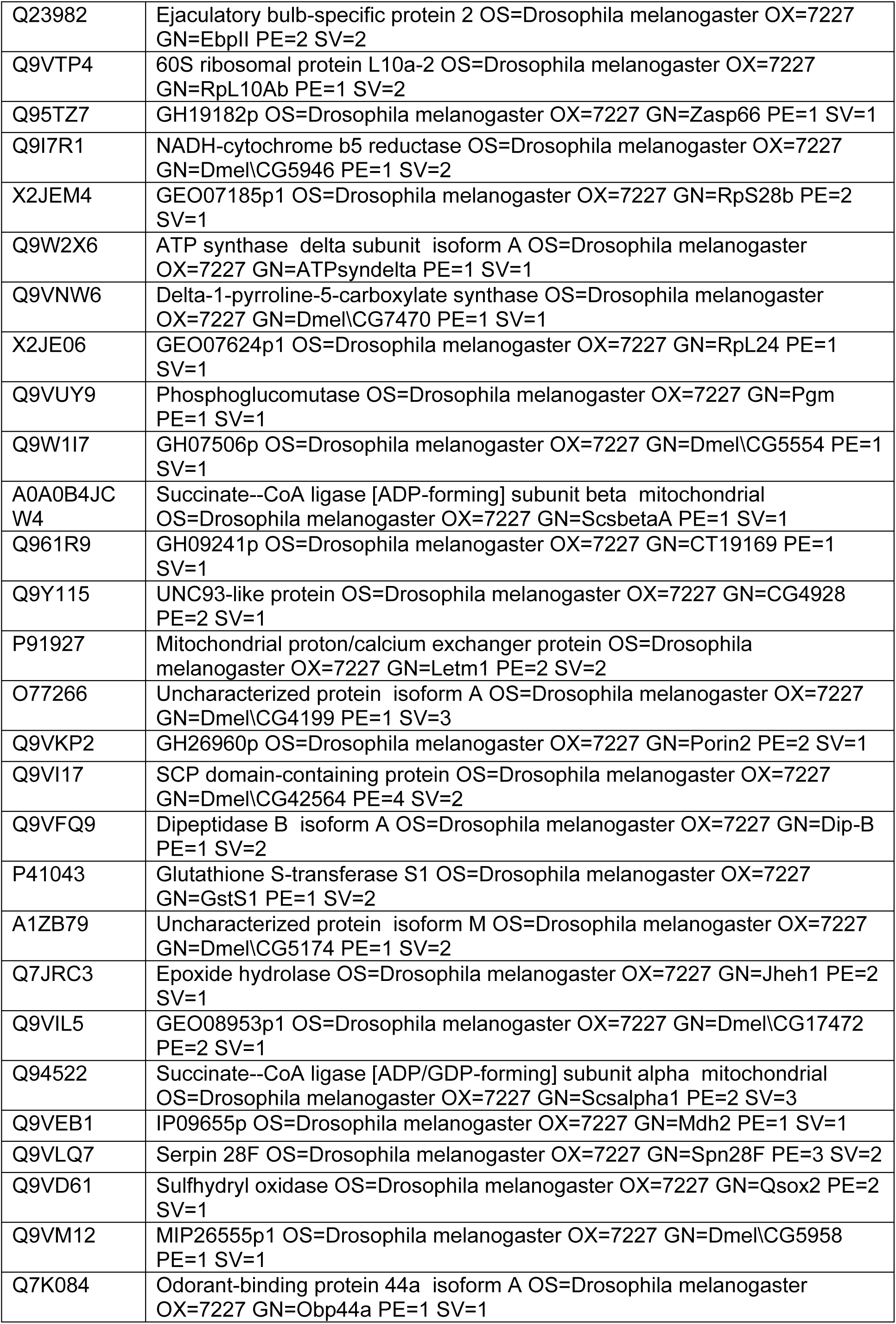
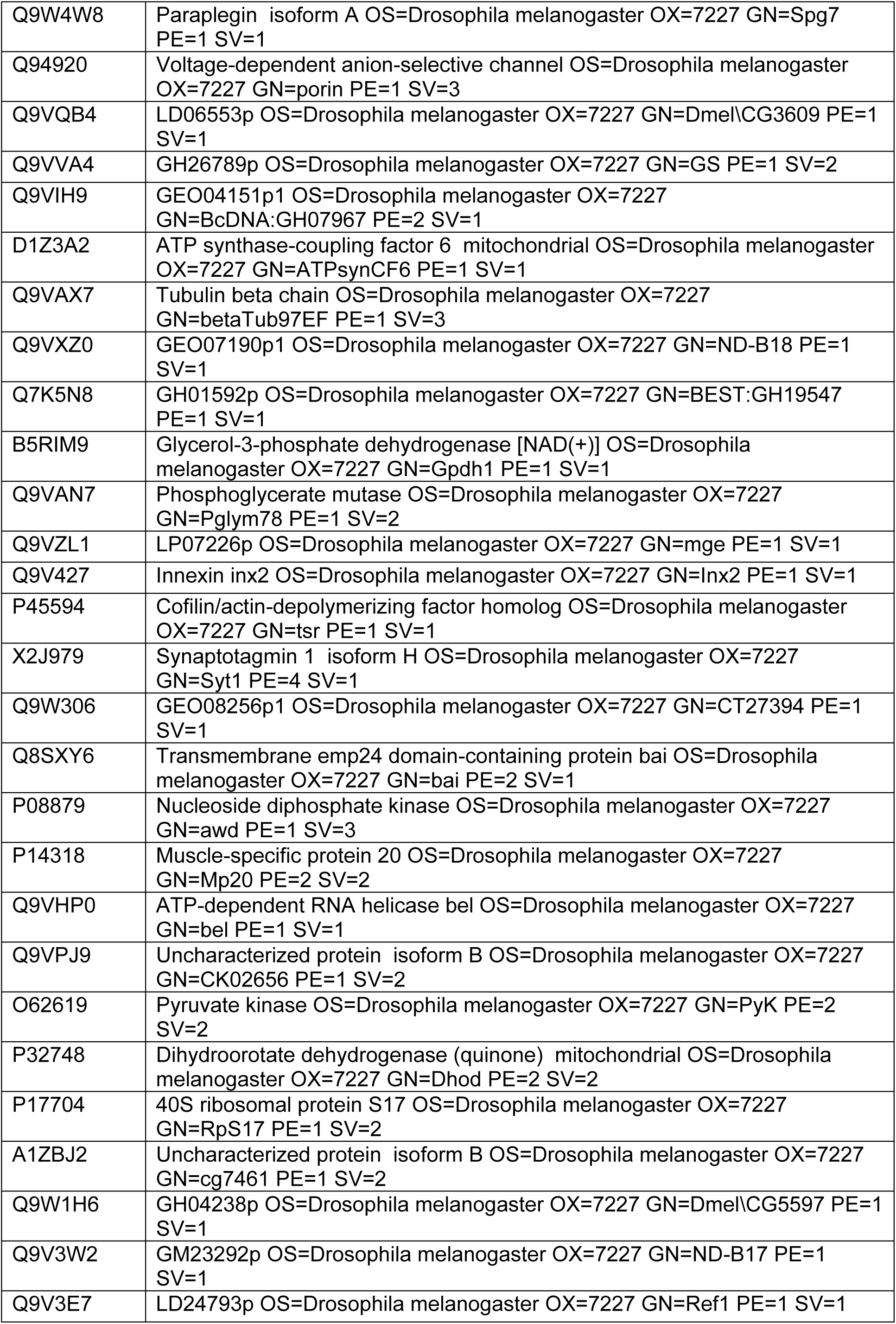
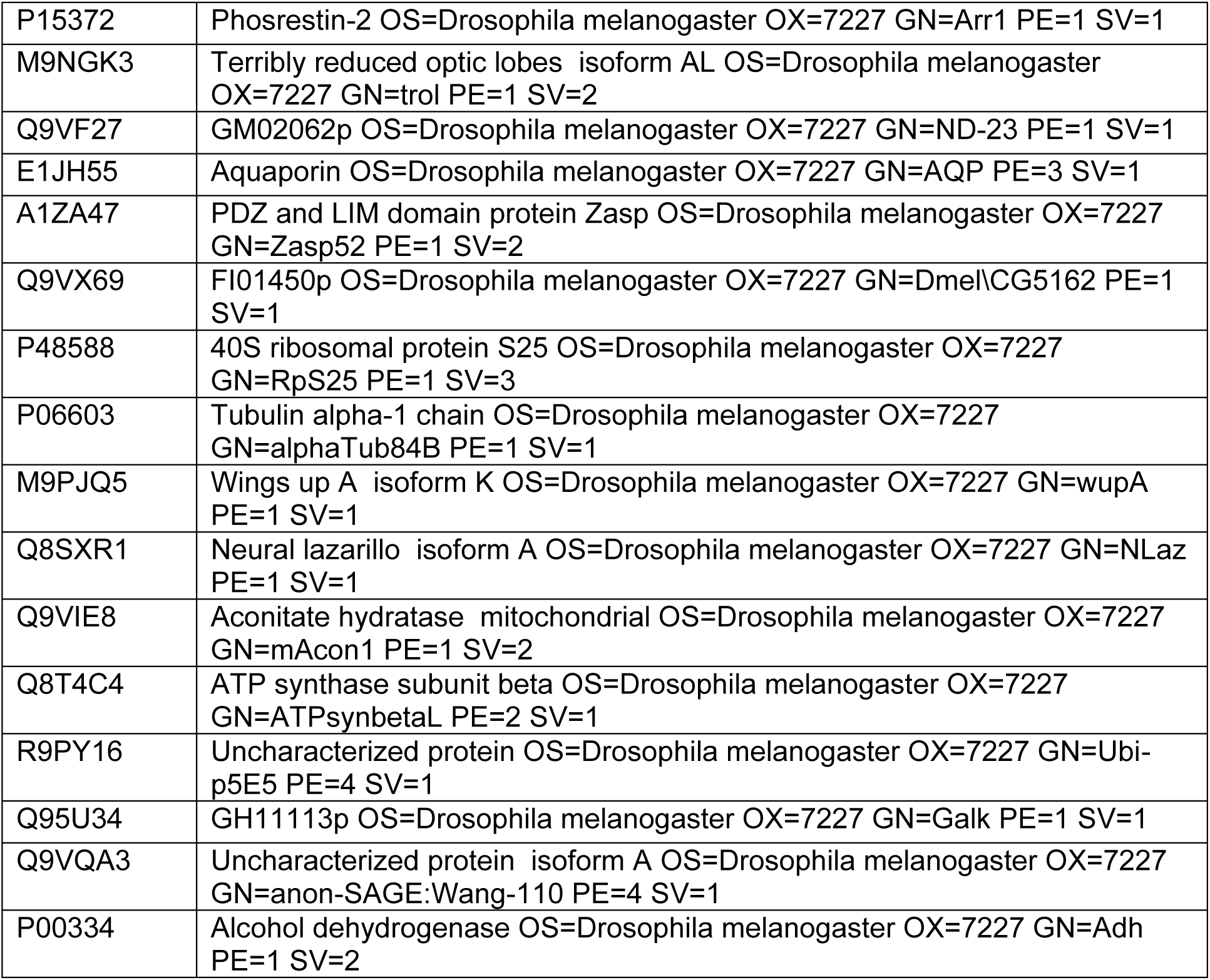
Label-free MS identified 516 proteins from the mitochondrial fractions of *Pink1^-^* and WT *D. melanogaster*

**Supplemental table 2.**
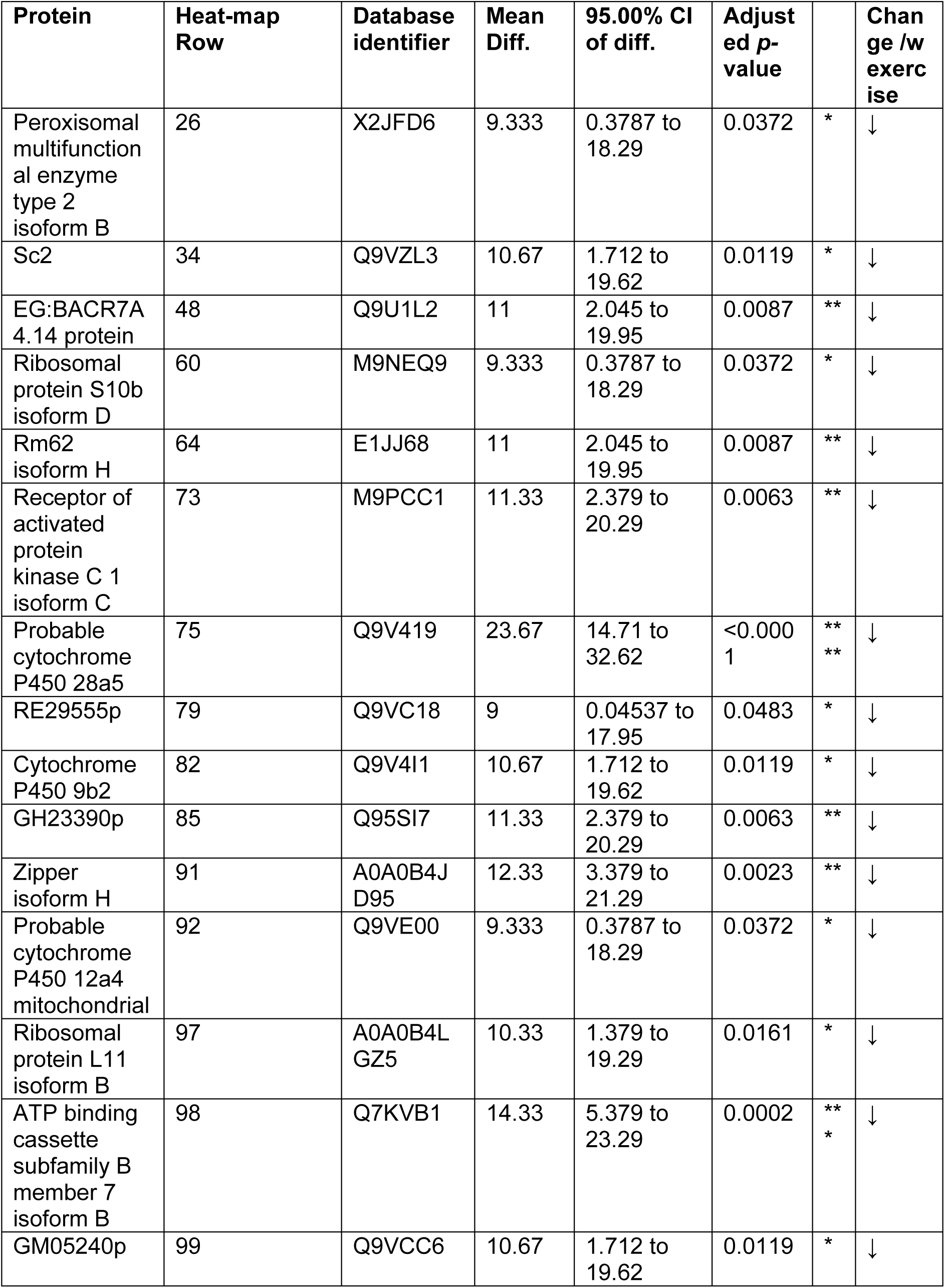
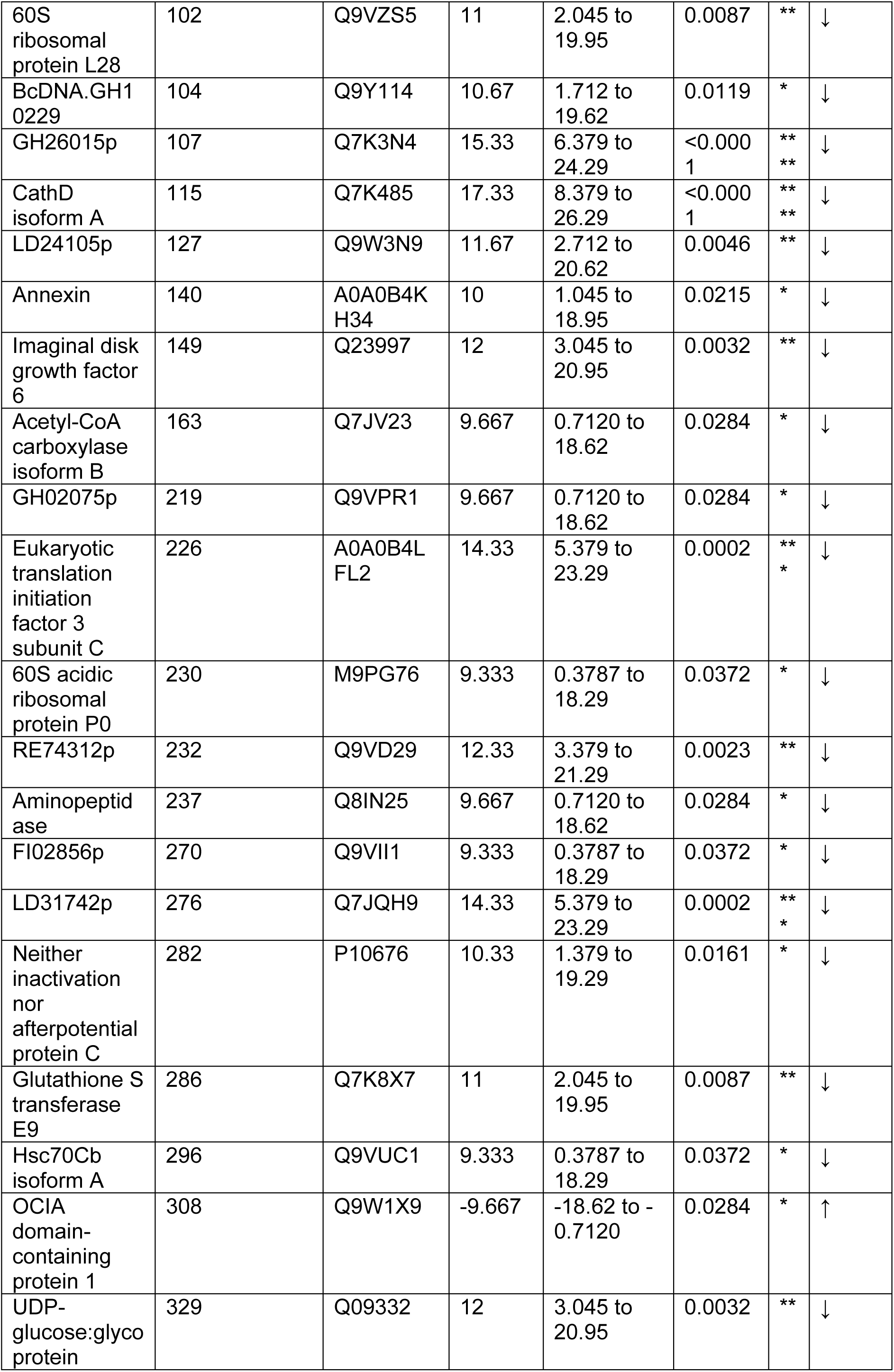
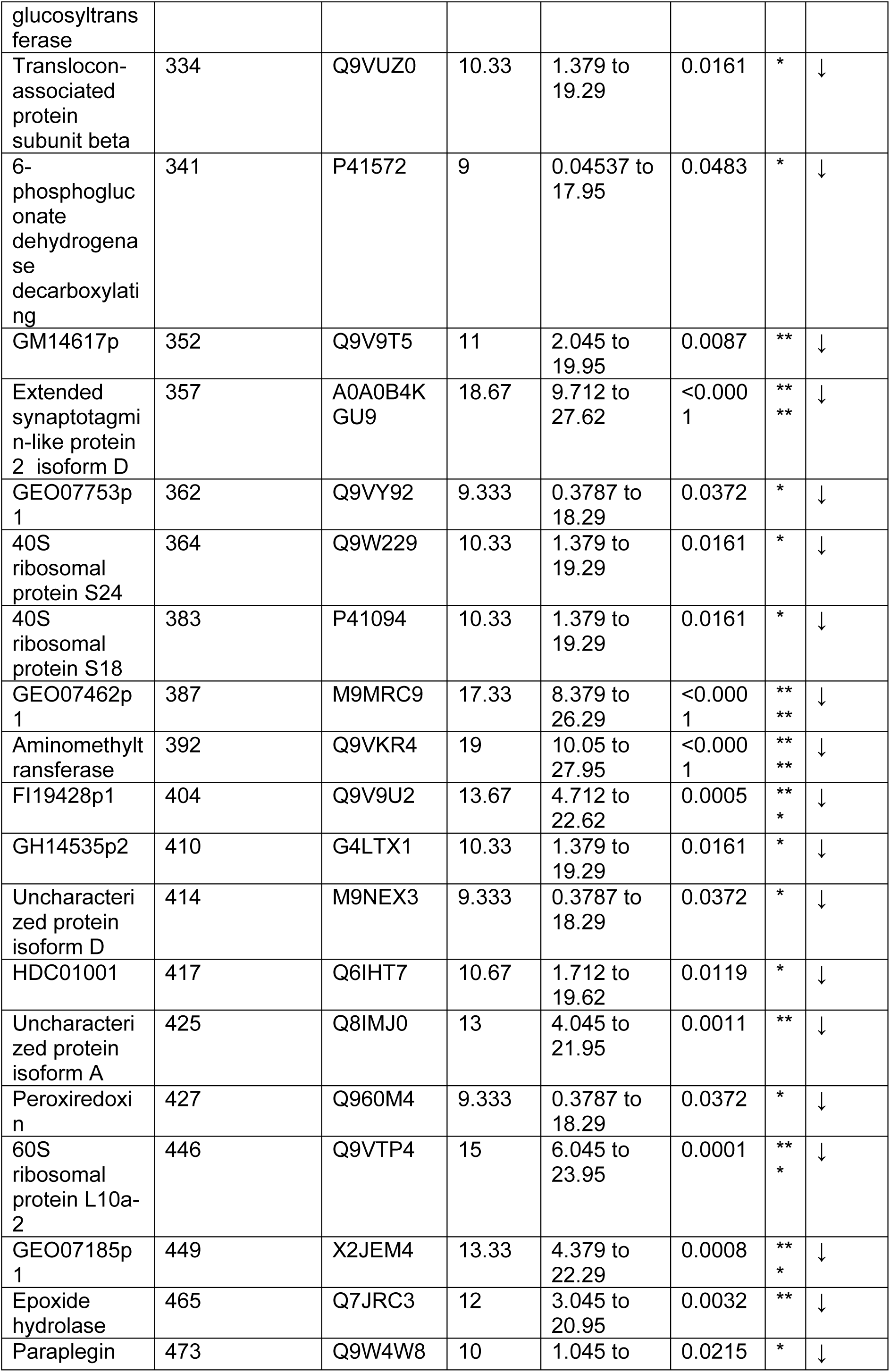
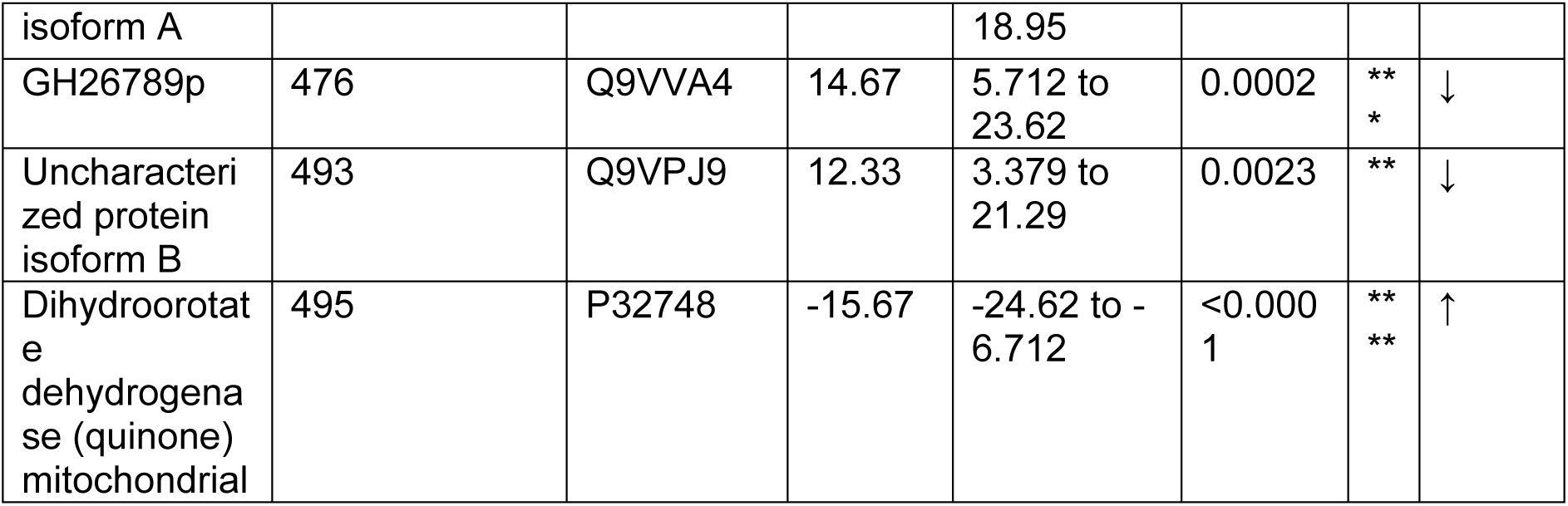
label-free MS identified 57 proteins with altered abundance between exercised and non-exercised *Pink1^-^* flies, 55 of which had decreased abundance.

**Supplemental table 3.**
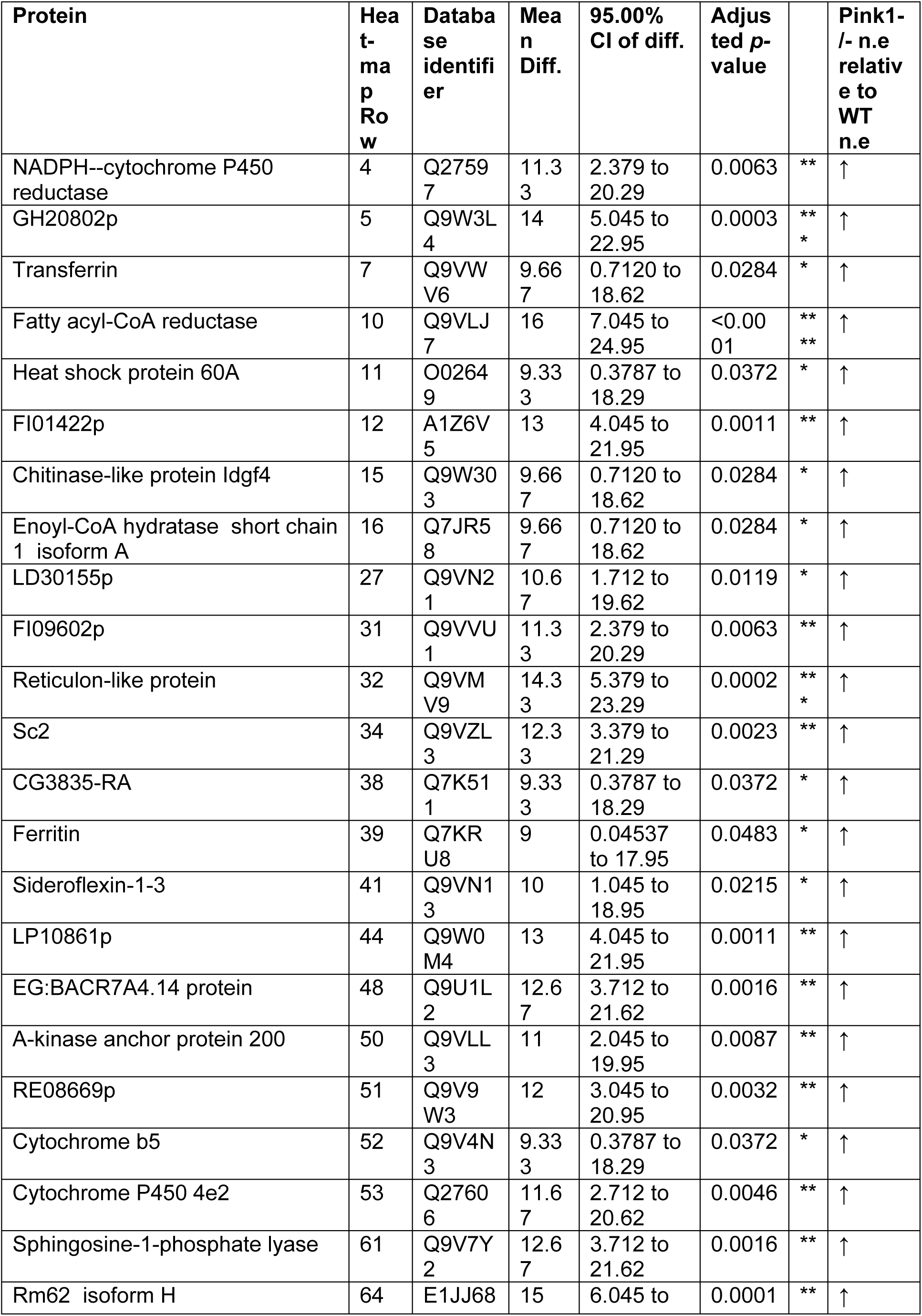
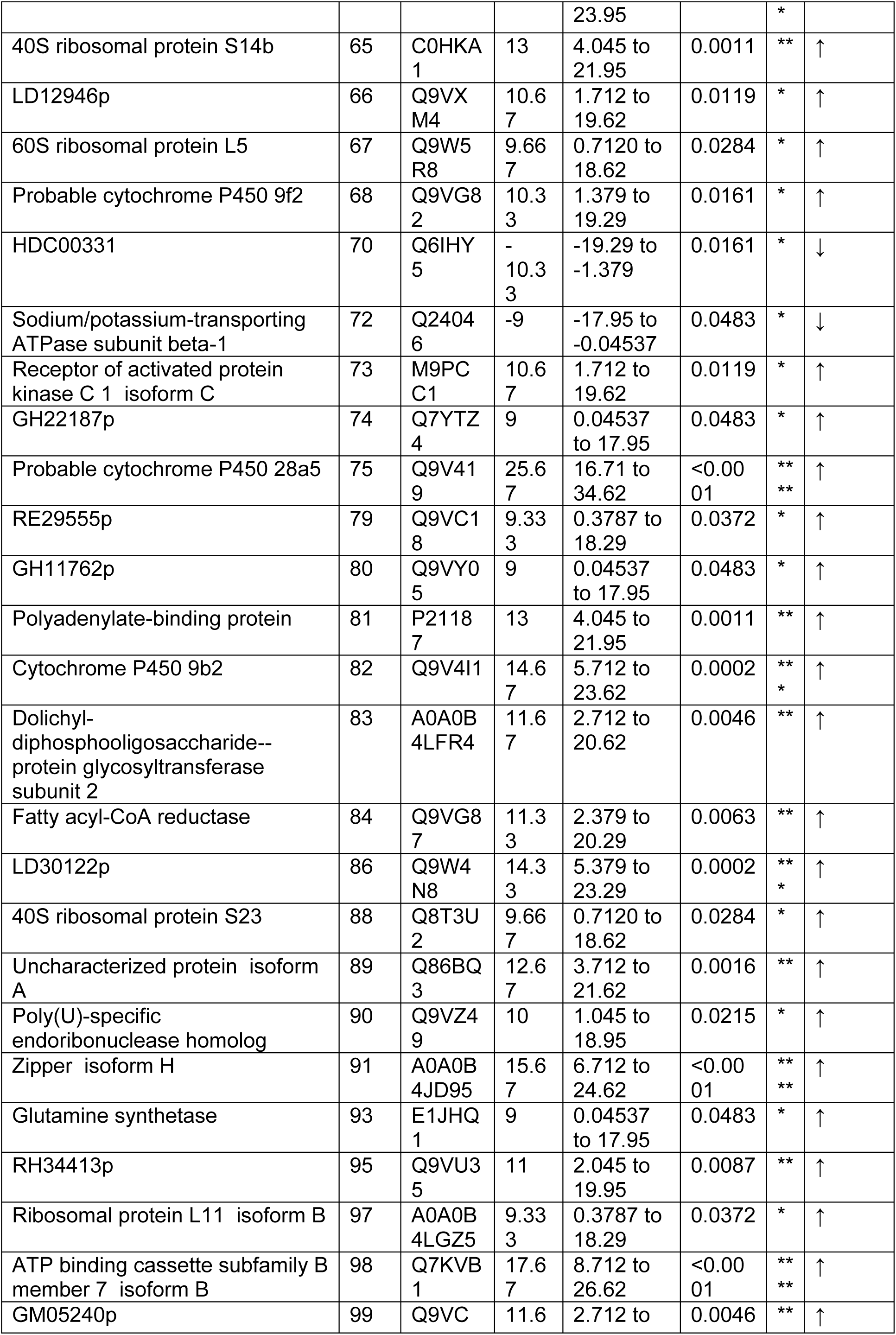
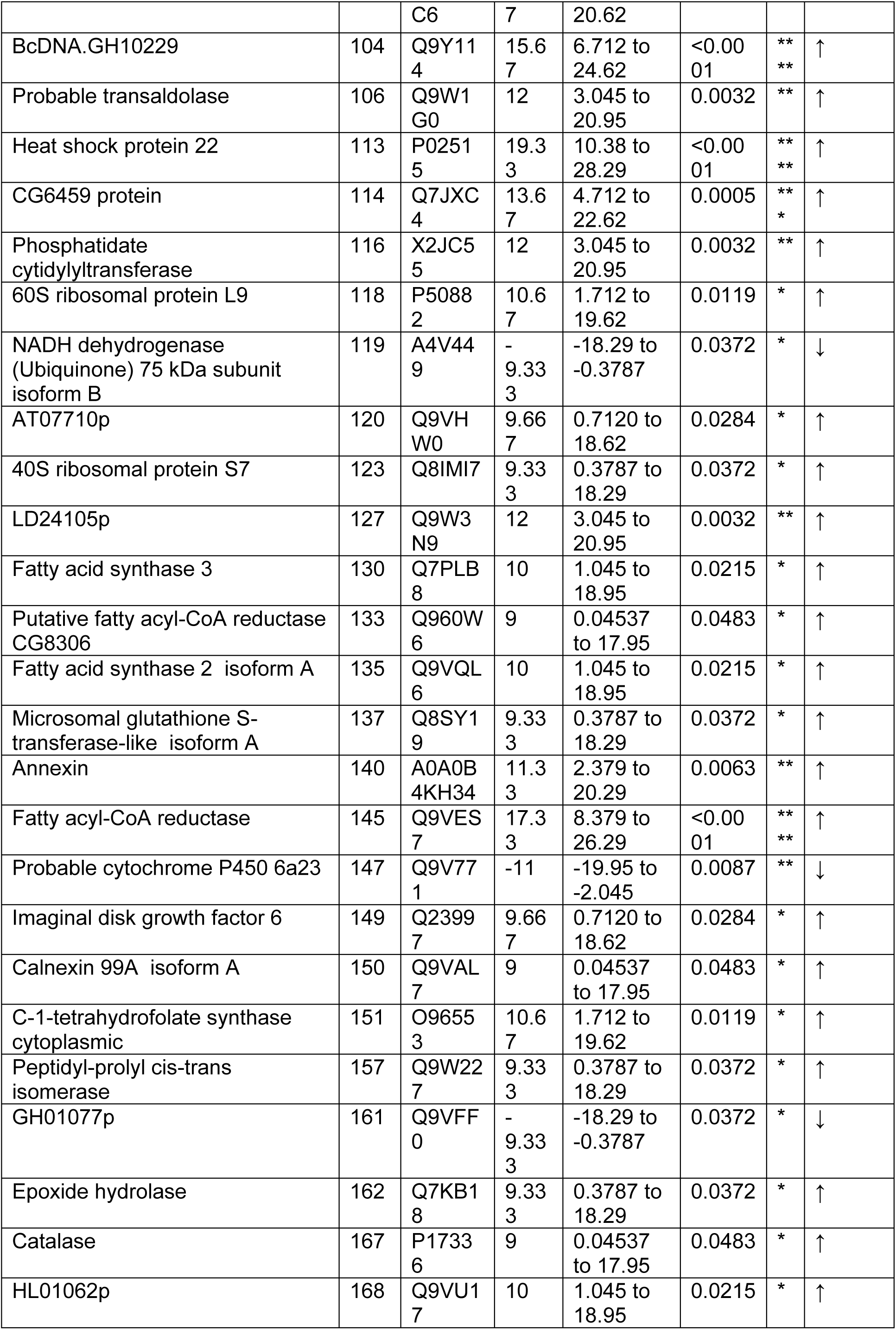
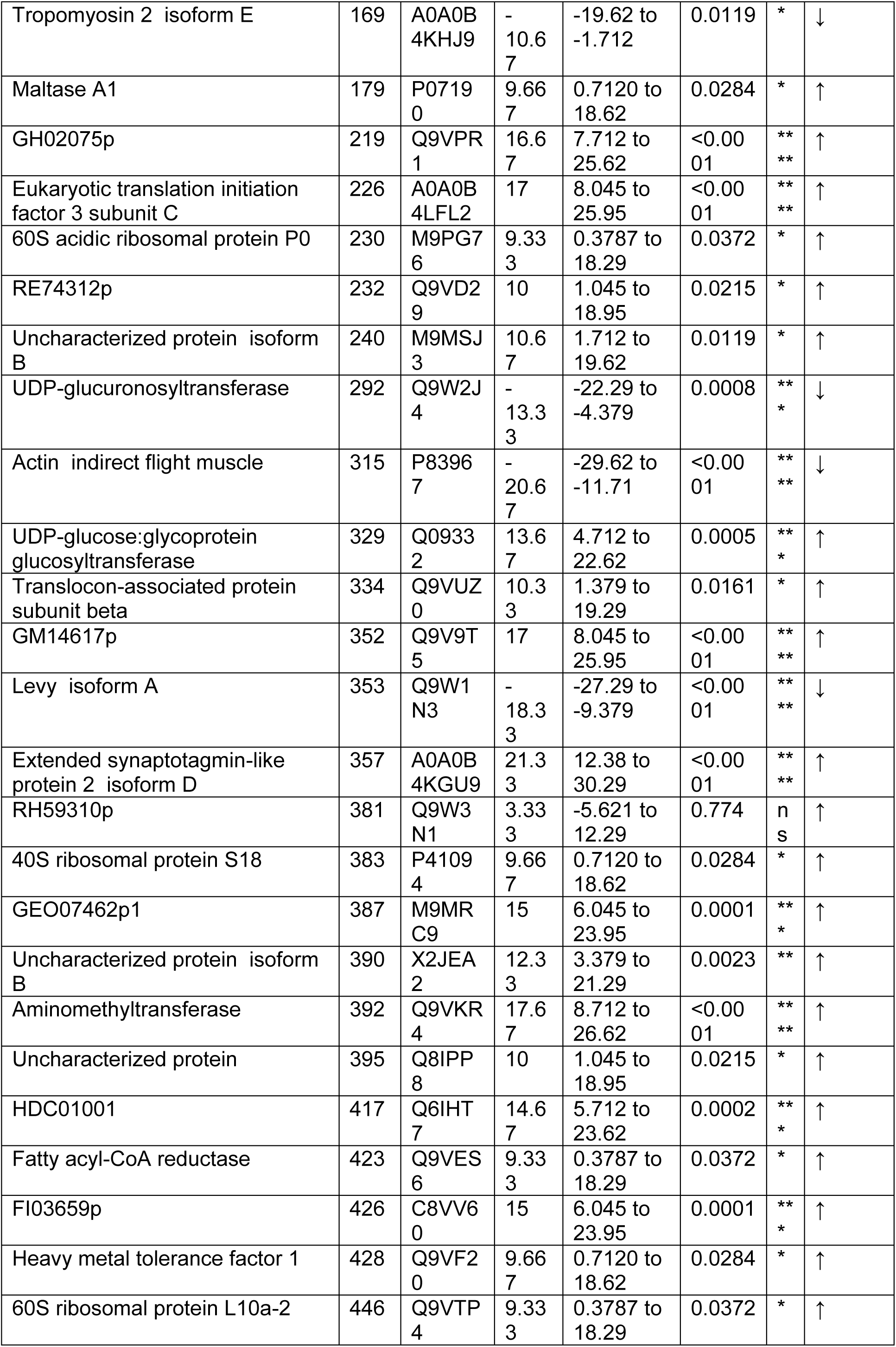
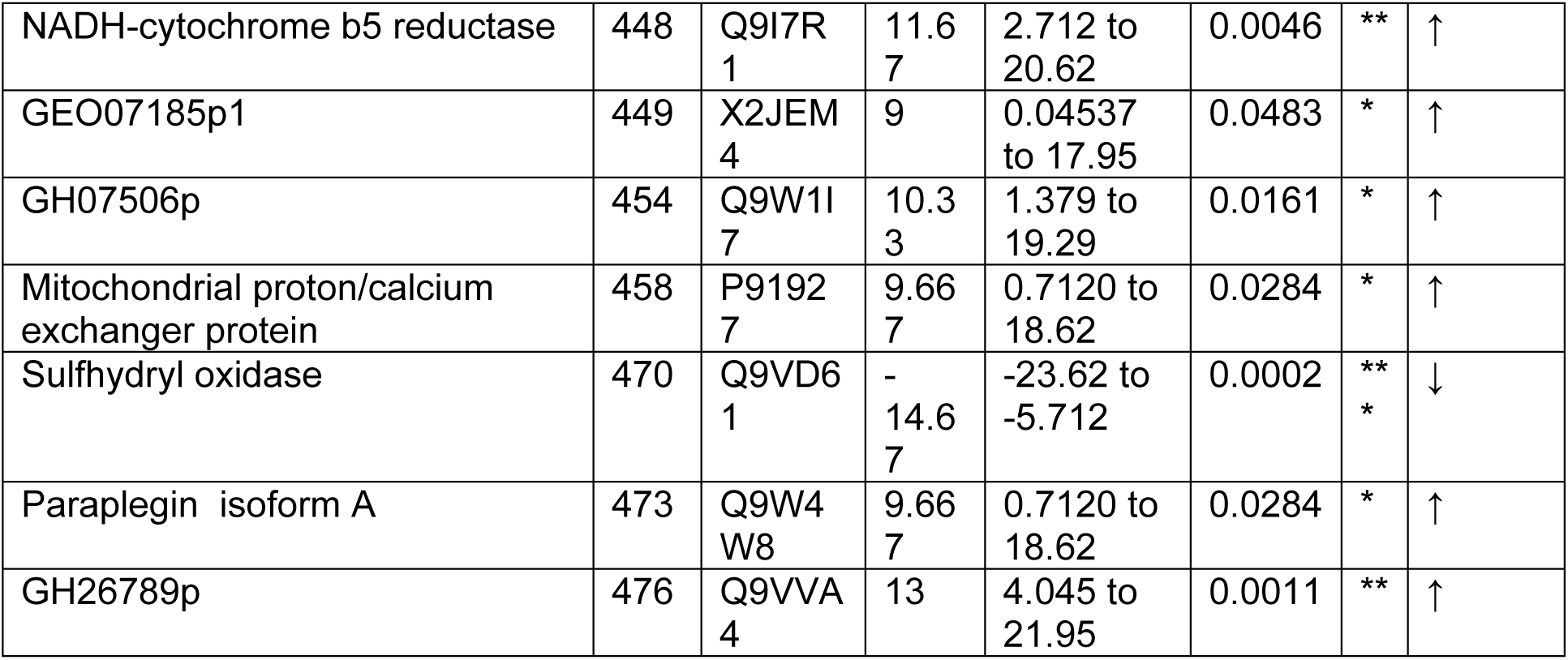
label-free MS identified 105 proteins with different abundance when comparing non-exercised *Pink1^-^* and WT flies

